# Marker-free characterization of single live circulating tumor cell full-length transcriptomes

**DOI:** 10.1101/2021.11.16.468747

**Authors:** Sarita Poonia, Anurag Goel, Smriti Chawla, Namrata Bhattacharya, Priyadarshini Rai, Yi Fang Lee, Yoon Sim Yap, Jay West, Ali Asgar Bhagat, Juhi Tayal, Anurag Mehta, Gaurav Ahuja, Angshul Majumdar, Naveen Ramalingam, Debarka Sengupta

## Abstract

The identification and characterization of circulating tumor cells (CTCs) are important for gaining insights into the biology of metastatic cancers, monitoring disease progression, and medical management of the disease. The limiting factor that hinders enrichment of purified CTC populations is their sparse availability, heterogeneity, and altered phenotypic traits relative to the tumor of origin. Intensive research both at the technical and molecular fronts led to the development of assays that ease CTC detection and identification from the peripheral blood. Most CTC detection methods use a mix of size selection, immune marker based white blood cells (WBC) depletion, and positive enrichment antibodies targeting tumor-associated antigens. However, the majority of these methods either miss out on atypical CTCs or suffer from WBC contamination. Single-cell RNA sequencing (scRNA-Seq) of CTCs provides a wealth of information about their tumors of origin as well as their fate and is a potent method of enabling unbiased identification of CTCs. We present unCTC, an R package for unbiased identification and characterization of CTCs from single-cell transcriptomic data. unCTC features many standard and novel computational and statistical modules for various analysis tasks. These include a novel method of scRNA-Seq clustering, named <u>D</u>eep <u>D</u>ictionary <u>L</u>earning using <u>K</u>-means clustering cost (DDLK), expression based copy number variation (CNV) inference, and combinatorial, marker-based verification of the malignant phenotypes. DDLK enables robust segregation of CTCs and WBCs in the pathway space, as opposed to the gene expression space. We validated the utility of unCTC on scRNA-Seq profiles of breast CTCs from six patients, captured and profiled using an integrated ClearCell^®^ FX and Polaris^TM^ workflow that works by the principles of size-based separation of CTCs and marker based WBC depletion.

## Introduction

Cancer ranks as a prime reason for death and a vital barrier to longer life expectancy in every country of the world (Sung *et al*., 2021). According to World Health Organization (WHO) estimates, in 2019 (Mathers, 2020) among 183 countries, cancer ranked as the first or second cause of death of people below the age of 70 years and ranked third or fourth in 23 countries (Sung *et al*., 2021). The primary reason for 90% of cancer-related deaths is metastasis (Bittner, Jiménez and Peyton, 2020), the process in which the cancer cells detach from the primary tumor, enter into the circulation, and eventually colonize distant organs, causing the spread of disease (Krebs *et al*., 2014; Siegel, Miller and Jemal, 2015). In order to metastasize, cancer cells secrete chemokines to attract immune cells (McAllister and Weinberg, 2014); (Liu and Cao, 2016), facilitating tumor proliferation and intravasation (Gajewski, Schreiber and Fu, 2013; Kitamura, 2018). After cancer cells enter the bloodstream, they are subjected to various stressors, including the lack of cell-cell and cell-matrix adhesion, shear pressures, and immune response. Despite this, a few cancer cells make it through the tortuous journey and leave the vasculature to a secondary site (Shenoy and Lu, 2016); (Follain *et al*., 2018).

Circulating tumor cells (CTCs) have recently attracted a lot of attention due to their critical role in tumor metastasis. Around 40% to 80% of patients with metastatic breast cancers have been found to have CTCs in their blood (Kwa and Esteva, 2018). The detection and characterization of CTCs obtained from patient blood offer clinically relevant insights into tumor metastasis and facilitate cancer diagnosis and treatment (Hong, Fang and Zhang, 2016). Various studies to date have unequivocally highlighted the association between the abundance of CTCs in peripheral blood and poor disease prognosis (Cristofanilli *et al*., 2004; Danila *et al*., 2007; Giuliano *et al*., 2011; Rack *et al*., 2014; Bork *et al*., 2015; Tsai *et al*., 2016). Epithelial to mesenchymal transition (EMT) is believed to play a crucial role in metastasis. Under EMT, tumor epithelial cells acquire mesenchymal-like features for easy entry into the bloodstream (Bulfoni *et al*., 2016).

Recently developed platforms for CTC capture rely on diverse principles. These include antibody-based capture (Nagrath *et al*., 2007; Riethdorf *et al*., 2007; Stott *et al*., 2010), size exclusion (Xu *et al*., 2015), immune cell depletion (Ozkumur *et al*., 2013), and dielectrophoresis (Chiu *et al*., 2016). CellSearch^®^, the only FDA-approved CTC capture platform, uses antibodies targeting the *EpCAM* (Epithelial cell adhesion molecule) antigen for capturing CTC from patients’ blood (Ignatiadis, Sotiriou and Pantel, 2012; Habli *et al*., 2020; Iyer *et al*., 2020). The expression of epithelial markers like *EpCAM* and *CK* (Creatine kinase) is used in affinity-based detection platforms to detect and count CTCs, but EMT arguably causes tumor cells to downregulate or lose expression of canonical epithelial markers, thereby making them hard to recognize and capture while in the circulation (Iyer *et al*., 2020). As such, marker-based enrichment strategies are sub-optimal for systematically charting heterogeneous CTC subpopulations (Miller, Doyle and Terstappen, 2010; Farace *et al*., 2011; Wang *et al*., 2016). Various CTC capture platforms based on biophysical characteristics of cancer cells have been established in recent years (Ferreira, Ramani and Jeffrey, 2016; Gabriel *et al*., 2016; Cheng *et al*., 2019). Negative selection for pan-leukocyte marker *CD45* has also been used as an alternative method. The promise of such antigen-agnostic platforms has not been explored adequately since the risk of immune cell contamination cannot be ruled out entirely (Ferreira, Ramani and Jeffrey, 2016; Gabriel *et al*., 2016). Isolation and molecular characterization of pure CTC populations by mRNA sequencing necessitates the development of precise analytic methods that are not based on epithelial markers and are able to spot the leukocyte contamination.

The advent of single-cell RNA sequencing (scRNA-Seq) has allowed in-depth, unsupervised analysis of CTC transcriptomes (Guo *et al*., 2015; Macosko *et al*., 2015; Kiselev *et al*., 2017; Butler *et al*., 2018; Wolf, Angerer and Theis, 2018; Kiselev, Andrews and Hemberg, 2019a; Chen *et al*., 2020; Ranjan *et al*., 2021). So far, most scRNA-Seq studies involving CTCs have used marker-based approaches to zero in on CTC subpopulations. Marker-agnostic methods for CTC annotation are rare and often incapable of confirming the malignant identity of the cells. The major challenges involved are as follows. 1. High levels of intra and inter-tumoural molecular diversity among malignant cells (Tirosh *et al*., 2016; Li *et al*., 2017). 2. Presence of CTCs in peripheral blood at an abysmally low concentration—one tumor cell within several millions of blood cells, even in patients with advanced metastatic disease (0–10 CTCs per mL of blood) (Alix-Panabières and Pantel, 2013). 3. CTCs often undergo EMT, thereby disguising their epithelial markers (Mikolajczyk *et al*., 2011); (Iyer *et al*., 2020). 4. Batch effect across scRNA-Seq studies (Kiselev, Andrews and Hemberg, 2019a); (Büttner *et al*., 2017).

To overcome these challenges, we present unCTC, an R package for unbiased characterization of CTC transcriptomes, in contrast with WBCs. unCTC features various standard and novel computational/statistical modules for clustering, copy number variation (CNV) inference, and marker-based characterization of CTC and non-CTC clusters obtained by analyzing the scRNA-Seq data. For clustering, DDLK, a deep dictionary learning based method is proposed. DDLK uses pathway scores at single-cell level to accurately segregate CTC and WBC populations. With unCTC, we demonstrated how *in silico* characterization of CTCs can unlock the power of marker-free CTC capture. For this we used the ClearCell^®^ Polaris™ workflow for size-based capture, immune cell depletion and single cell gene expression profiles of potential CTCs (Warkiani *et al*., 2014; Ramalingam *et al*., 2017). The unCTC workflow confers phenotypic identity on the captured cells through multi-factorial analyses of the single cell expression profiles.

For validation of unCTC, we used seven distinct single-cell RNA-Seq (scRNA-Seq) datasets of single CTCs and WBCs (Aceto *et al*., 2014; Ting *et al*., 2014; Yu *et al*., 2014; Sarioglu *et al*., 2015; Jordan *et al*., 2016; Velten *et al*., 2017; Zheng *et al*., 2017). unCTC aided integrative analysis perfectly segregated CTCs and WBCs. Apart from this, we subjected ClearCell Polaris selected potential CTC full-length transcriptomes obtained from six individuals with breast cancer of three molecular subtypes (ER-/PR-/HER2-, ER+/PR+/HER2-, and ER-/PR-/HER2+) to the unCTC analysis pipeline. As control we used independent scRNA-Seq data of breast CTCs and WBCs. Our unCTC based analysis of the data confirmed CTC capture efficiency of the ClearCell-Polaris microfluidic workflow for marker-free CTC capture.

## Results and Discussion

### Overview of the unCTC workflow

Identification and characterization of CTC using scRNA-Seq profiles is ever-challenging due to the dynamic nature of CTC phenotype. The unCTC workflow features a number of methods that help in unbiased identification and characterization of single CTC transcriptomes. Clustering of scRNA-Seq profiles is an important step towards this. Here we present a robust approach for clustering single cell transcriptomes in a metaspace, spanning pathways, whose enrichment scores are computed on single-cell gene expression readouts. Single-cell expression data are typically sparse (Kiselev, Andrews and Hemberg, 2019b; Tian *et al*., 2019). Pathway scores computed on gene-sets alleviate this problem to a great extent (Li *et al*., 2017; Chawla *et al*., 2021), thereby assisting in robust detection of cellular subtypes. For unsupervised clustering, each of the normalized and log-transformed expression vectors associated with CTCs is converted into a vector of pathway enrichment scores, calculated using GSVA (gene set variation analysis) (Hänzelmann, Castelo and Guinney, 2013). Such a transformation neutralizes batch effects (Kim *et al*., 2018), and unravels cellular heterogeneity from a rather functional/mechanistic point of view (H. Ding *et al*., 2019; Ramirez *et al*., 2020; Wang *et al*., 2020). DDLK incorporates the K-means clustering cost into the Deep Dictionary Learning (DDL) framework. Shallow learning and data dependency are the main caveats of dictionary learning and deep learning respectively. Deep dictionary learning aims to mitigate these challenges (Tariyal *et al*., 2016). DDLK is an example of semi-supervised clustering which projects the single-cell gene expression data onto a range of well-understood biological pathways to obtain robust cellular clusters.

While DDLK robustly identifies phenotypically similar cell-groups from scRNA-Seq data containing CTC expression profiles, cluster annotation may still remain elusive. To address this, we integrated inferCNV function into the unCTC R package. InferCNV is an existing method, capable of inferring copy number variation (CNV) from single-cell gene expression data (Couturier *et al*., 2020). Chromosomes in cancer cells undergo substantial aberration. InferCNV has been proved to be useful in capturing approximate CNV locations at single-cell resolution (Durante *et al*., 2020; Zhou *et al*., 2020). InferCNV works as a sounding board for cell type characterization, especially zeroing in on the malignant origin of CTCs. Further, inferCNV along with cytoband information based on GRCH37 (Barrios and Prieto, 2017) also pinpoints the precise position of chromosomal aberrations at the level of chromosomal arms, aiding in the identification of altered genes.

CTCs undergo EMT and other biophysical stress during their journey to distant organs. In this process, they partially lose their epithelial phenotype. Univariate differential expression studies may turn out to be limitedly helpful in such scenarios. To circumvent this, unCTC allows cumulative measurement of enrichment of a range of canonical markers indicating malignant/epithelial/immune origin with the help of Stouffer’s method (Stouffer *et al*., 1949). In our hands such gene set-based approaches turn out to be fruitful in bolstering single marker-based and inferred CNV based characterization of cell-groups. The complete unCTC workflow is outlined in **Figure 1**.

**Figure 1.**
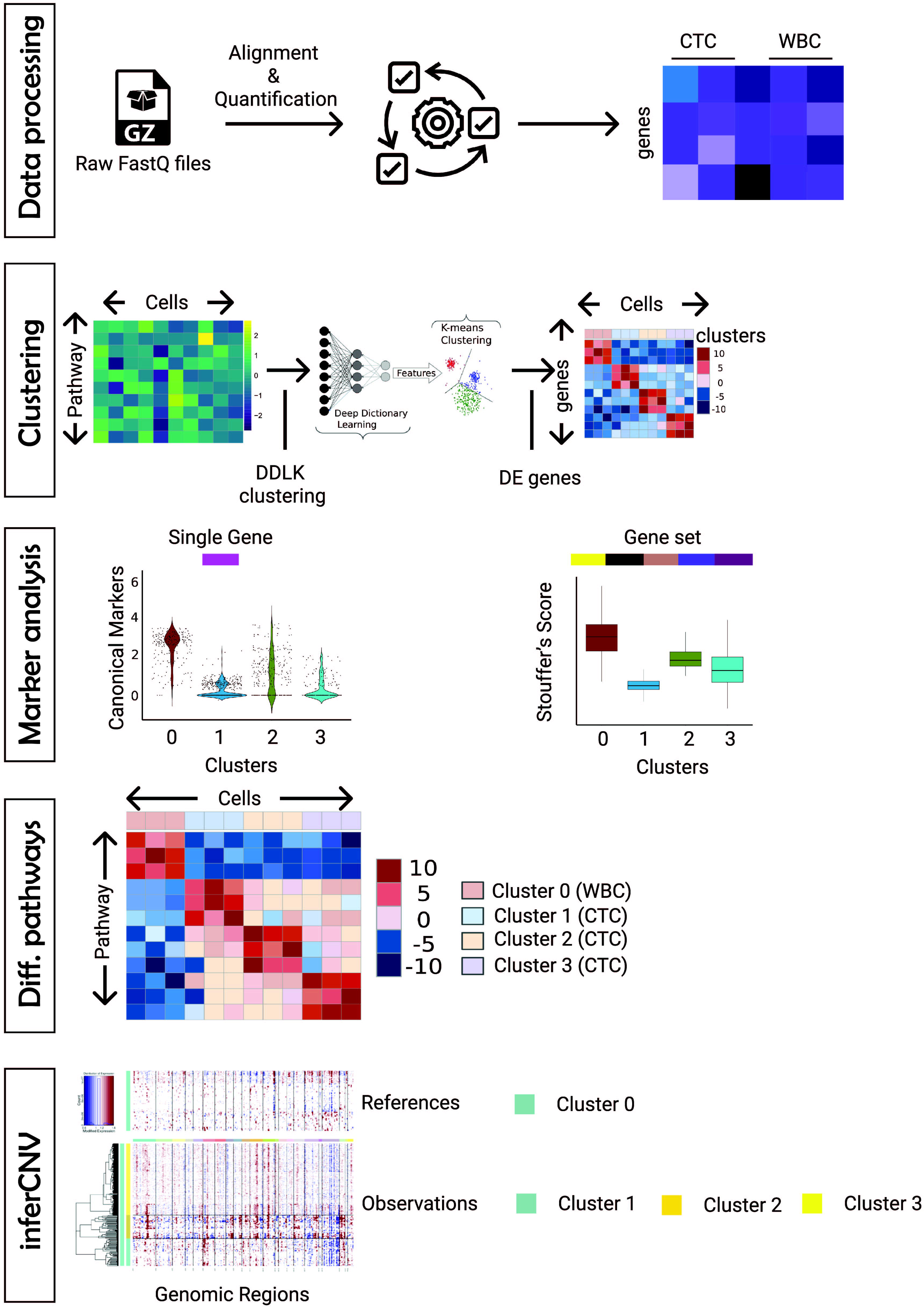
unCTC: A unified, end-to-end computational framework for marker-free characterisation of CTCs. Schematic diagram depicting the analysis workflow as well as the key methods supported by the unCTC R package. The first step involves processing raw FASTQ files to obtain the expression matrix. Novel DDLK clustering method is used to robustly cluster single CTC transcriptomes. Notably, DDLK works on pathway enrichment scores as opposed to expression values. Clusterwise differential expression analysis is performed to gain insights into diverse CTC and WBC subtypes. Expression levels of well-known epithelial and immune markers are tracked to approximate broad cell-type identities. Similar analysis is also carried out at the level of well-known gene-sets/pathways. Further, differential enrichment of pathway specific genes can be analysed to infer functional attributes. Finally, expression based pseudo-CNV inference allows unbiased characterisation of the identified clusters, thereby highlighting the malignant cells.

### Marker-free capture of CTCs

Breast cancer is the most frequent type of cancer and one of the top causes of cancer-related mortality (Kamal *et al*., 2017). In 2020, breast cancer overtook lung cancer as the world’s most common cancer. About 90% of breast malignancy related fatalities are attributable to metastasis (Zhang *et al*., 2021). Breast cancer appears to be the form of cancer in which CTCs have been studied the most (Bidard, Proudhon and Pierga, 2016). The expression of epithelial markers, including *EpCAM* and *CK*, has traditionally been used in affinity-dependent detection platforms to recognize and count CTCs, but these markers are downregulated during EMT. Furthermore, most fluorescence-activated cell-sorting techniques face challenges due to acute scarcity of CTCs, where the scarcity is typically less than 1 CTC/10 mL of blood in nonmetastatic malignancies (Thery *et al*., 2019). Due to this inadequacy, a marker-free, robust method is necessary for detecting and enriching CTCs in a large pool of blood cells. Marker-free methods for isolating CTCs are appealing because they enable researchers to examine a greater number of CTCs that would otherwise be missed due to variable or absent protein (label) marker expression on the CTC surfaces. It was possible to establish a marker-free method for isolating CTCs by integrating the ClearCell^®^ FX and Polaris^TM^ systems (Iyer *et al*., 2020). As part of this, CTCs are enriched in two steps - size-based enrichment by ClearCell, followed by *CD45* (leukocyte marker) and *CD31* (endothelial cell marker) based negative depletion by Polaris (Warkiani *et al*., 2014; Ramalingam *et al*., 2017) **(Figure 2)**. Using the ClearCell FX and Polaris systems, we collected 81 single CTCs from six women with breast cancer of three subtypes (ER-/PR-/HER2-, ER+/PR+/HER2-, and ER-/PR-/HER2+) **(Supplementary Table S1)**. 72 CTCs finally qualified the quality control criteria **(Supplementary Table S2)**. In the subsequent sections we illustrate unCTC based characterization of these cells, in contrast to two other best practice integrative analysis methods namely — Seurat (Hao *et al*., 2021) and Symphony (Kang *et al*., 2020).

**Figure 2.**
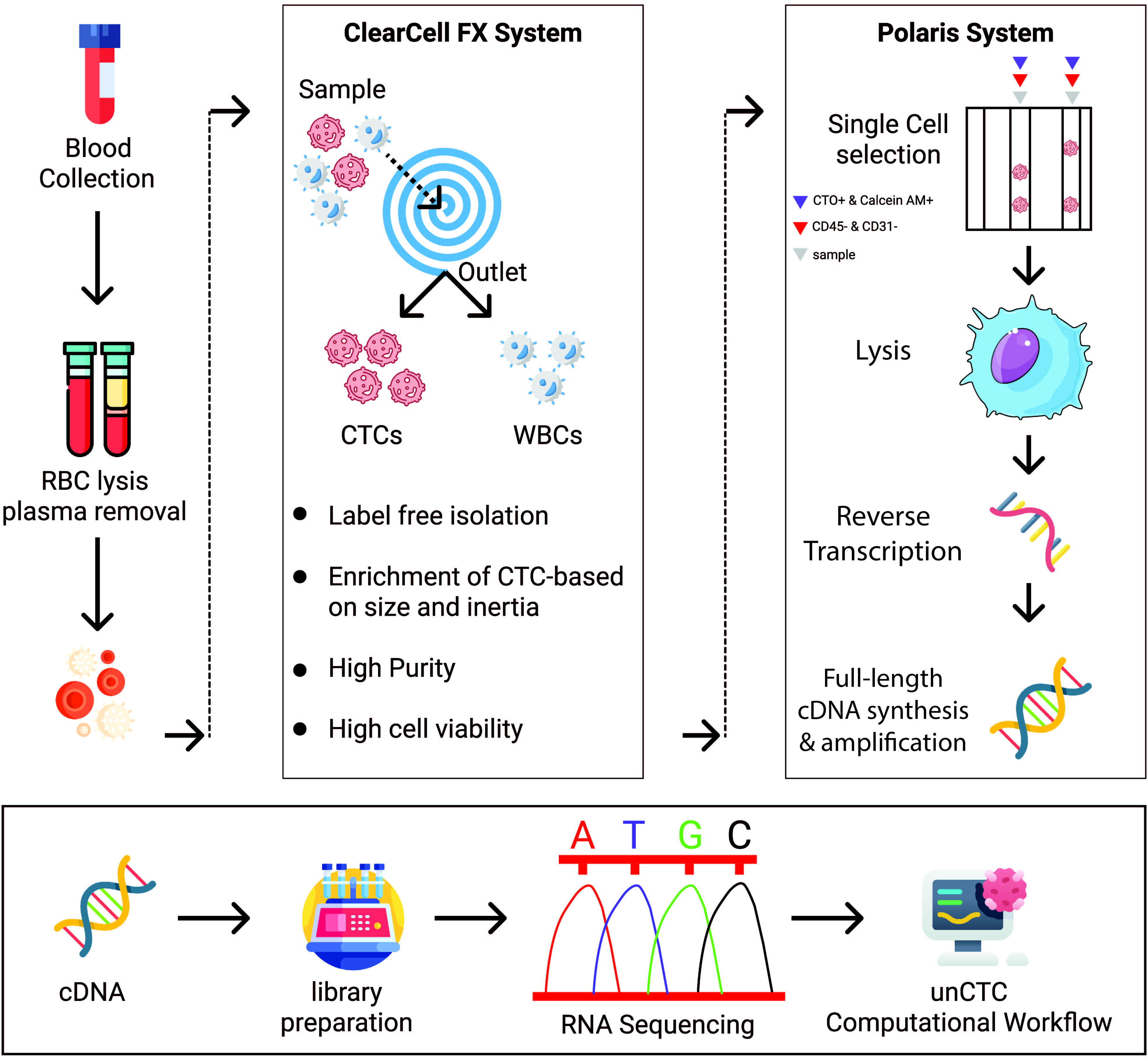
ClearCell^®^ FX and Polaris^TM^ workflow for marker-free enrichment of CTCs. (A) The schematic diagram depicting the key steps involved in the capture and isolation of CTCs using a two pronged system. ClearCell^®^ FX uses a spiral chip to size-sort CTCs. Polaris^TM^ performs single cell capture and cDNA synthesis of potential CTCs after depletion of cells that are CD45/CD31 positive. Finally cDNA thus received is subjected to library preparation and RNA-sequencing.

### DDLK clustering leads to near perfect segregation of CTCs and WBCs

The main aim of unCTC is to enable segregation of the CTC and WBC populations, obtained after unbiased microfluidic enrichment of CTCs in patient blood. This problem is fundamentally different from identifying CTC clusters or deciphering functional heterogeneity among single-cell transcriptomes, a method that typically requires unsupervised clustering of expression vectors. To cater to this objective, we chose to project expression vectors in a meta-space spanned by well-characterized biomolecular pathways using GSVA that when supplied with selected pathways, convert given expression vectors into vectors comprising pathway enrichment scores (Hänzelmann, Castelo and Guinney, 2013). This is particularly advantageous and confers robustness in data integration tasks (Jin *et al*., 2014). Expression vectors, after conversion into vectors of pathway enrichment scores, are used for clustering the associated single cell transcriptomes. Existing deep learning-based clustering techniques use stacked autoencoders (Peng *et al*., 2016; Xie, Girshick and Farhadi, 2016; Yang *et al*., 2017; Fard, Thonet and Gaussier, 2020) or their convolutional counterparts (Guo *et al*., 2017; Yang *et al*., 2019). Unlike deep dictionary learning (DDL) (Tariyal *et al*., 2016), the problem with autoencoders is that they have to estimate the parameters (encoder network + decoder network) twice. This leads to overfitting and general degradation of results. Prior studies have shown DDL to be the go-to framework for data constrained scenarios (Mahdizadehaghdam *et al*., 2019) (Tang *et al*., 2020) (Fu *et al*., 2019) instead of conventional deep learning. DDLK (clustering technique incorporated in unCTC) incorporates K-means clustering cost into the deep dictionary learning (DDL) framework, thereby enabling clustering of data of all sizes. We compared unCTC to two popular single-cell integrative analysis suites, Seurat (Butler *et al*., 2018) and Symphony (Kang *et al*., 2020). Seurat returns unsupervised clusters, whereas Symphony returns integrated single cell representation alone.

Presently most single-cell studies involve integration of scRNA-Seq datasets coming from different biological replicates, giving rise to significant batch effects (Sinha *et al*., 2019). Also, due to small amounts of starting RNA, single-cell data, even if it comes from a single chip, exhibits cell-to-cell technical variability. It is, therefore, imperative to ensure that a single-cell pipeline is robust to such variance factors. To validate this, we constructed a challenging multi-study (141 CTCs spanning breast, lung and pancreatic cancers; 1037 WBCs) dataset (Aceto *et al*., 2014; Ting *et al*., 2014; Yu *et al*., 2014; Sarioglu *et al*., 2015; Jordan *et al*., 2016; Velten *et al*., 2017; Zheng *et al*., 2017) for comparative assessment of unCTC in contrast to Seurat and Symphony **(Supplementary Table S3)**. We subjected the dataset to unCTC and two other best practice methods Seurat and Symphony. Symphony and unCTC managed to visually segregate CTCs and WBCs, whereas Seurat failed to separate the two categories **(Figure 3)**. Symphony does not cluster the cell. unCTC was able to find CTCs as part of a single cluster **(Figure 4C,D)**.On the contrary, clusters returned by Seurat had a mixture of both cell types **(Figure 4A, B)**.

**Figure 3.**
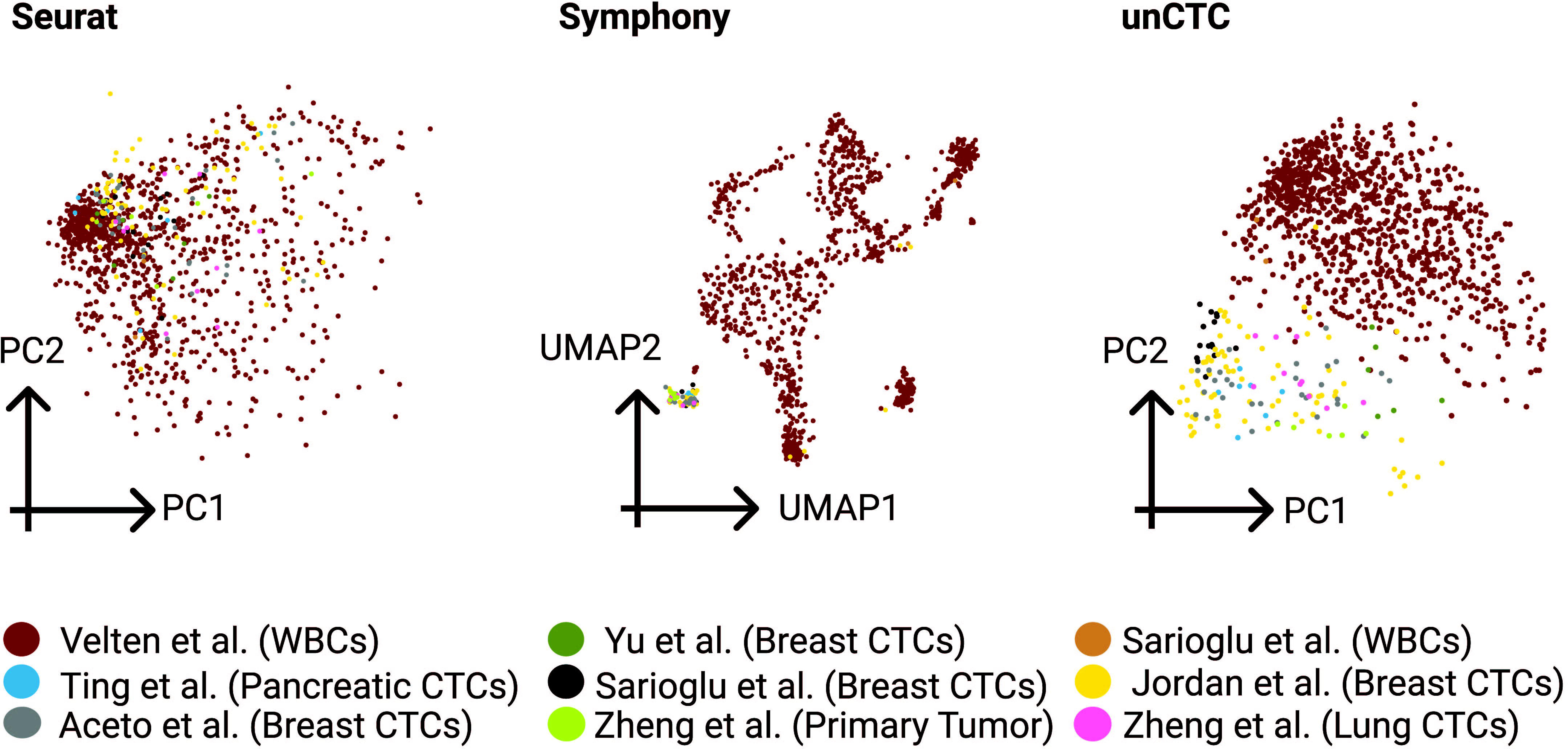
unCTC enables integrative analysis of CTCs and WBCs. A multi-study CTC/WBC dataset was created to test the efficacy of unCTC alongside two best practice integrative scRNA-Seq analysis pipelines namely Seurat and Symphony. Both Symphony and unCTC accurately segregated the CTCs and WBCs, as visible from the low-dimensional representation of the cells.

**Figure 4.**
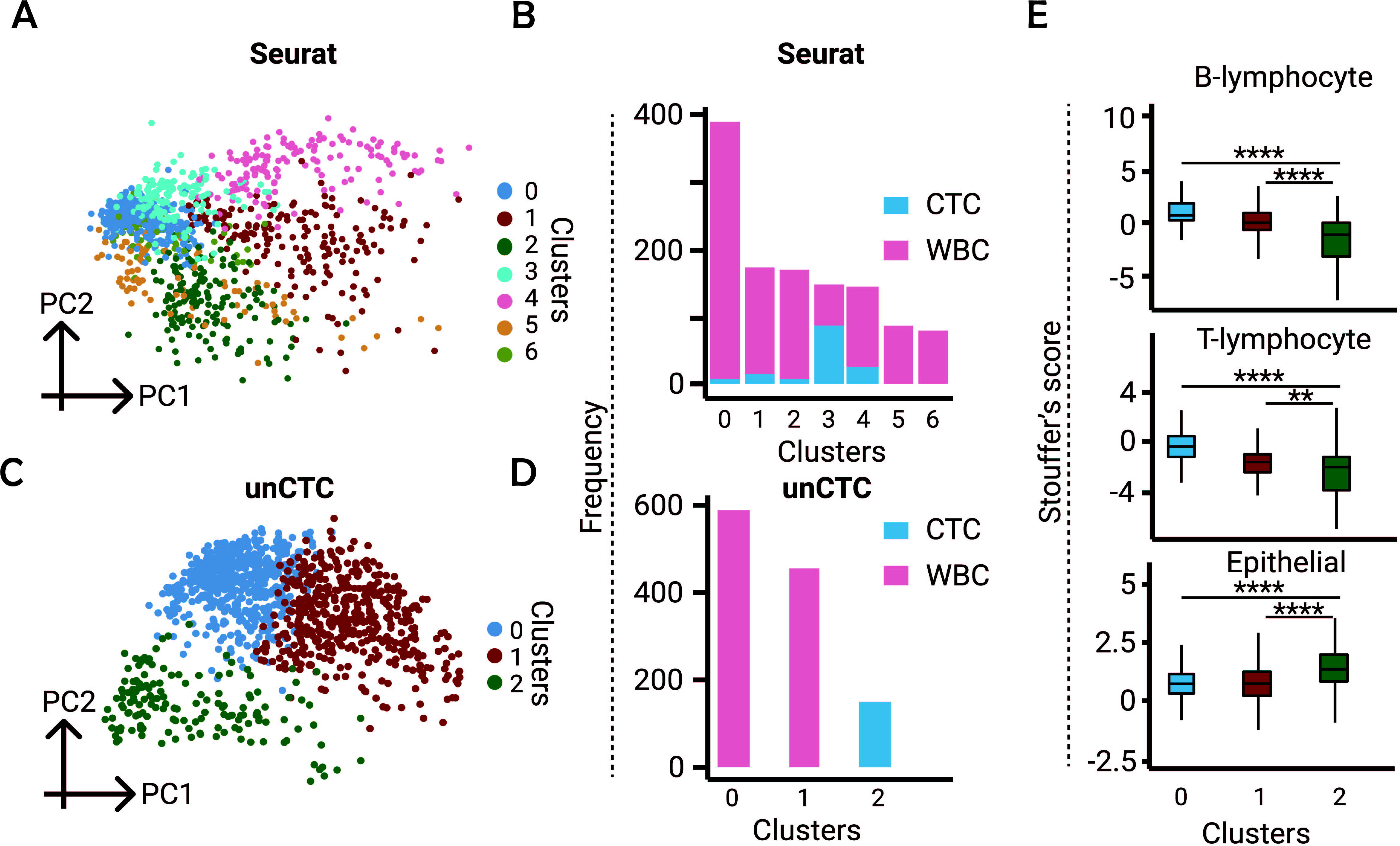
Cluster purity. (A) Visualisation of cluster identity using Seurat. (B) CTC/WBC composition for each cluster, obtained using Seurat. (C,D) Equivalent figures depicting unCTC clustering and cluster purity respectively. (E) Boxplots depicting distribution of Stouffer’s scores computed basis known B cell, T cell and epithelial cell markers respectively, for cells in each of the unCTC identified clusters.

While clustering is the most popular means for unsupervised multivariate analysis of single cells, cell-lineage annotation requires examination of marker-gene expression. Typically, univariate statistics are applied to user-selected canonical markers to confer lineage identity on CTCs and WBCs. Given the unpredictable expression dynamics of single markers in CTCs, it is often useful to measure combined upregulation of tens of markers, reducing dependency on individual genes. To this end, we used Stouffer’s method to combine expression levels of a range of markers associated with cell-lineages of interest **(Supplementary Table S4)**. **Figure 4E** highlights cluster-specific enrichment scores of genes related to B and T lymphocytes, as well as epithelial markers. Out of the three clusters retrieved by unCTC, cluster 2 exhibited the highest enrichment of epithelial markers. This finding is concordant with annotations sourced from the studies.

### unCTC recognizes CTCs selected by the ClearCell FX and Polaris workflow

We have recently demonstrated CTC characterization using supervised machine learning methods (Iyer *et al*., 2020). Given the dynamic nature of CTC phenotypes, it is however useful to characterize single CTC transcriptomes by unsupervised means. Further, classification-based characterisation approaches are fallible in scenarios where the obtained CTCs are of atypical phenotypes. unCTC alleviates this shortcoming by bringing to bear a spectrum of unbiased single-cell characterization tools. As an extended validation, we subjected the 72 filtered single cell transcriptomes associated with potential CTCs captured by the ClearCell^®^ FX and Polaris^TM^ workflow. These come from a total of six women entailing three major subtypes of breast cancer — ER-/PR-/HER2-, ER+/PR+/HER2-, and ER-/PR-/HER2+. As control we also considered the CTC dataset published by Ebright et al. (Ebright *et al*., 2020) that comprises 824 cells from 45 patients with breast cancer of the ER+/PR+/HER2-subtype. For WBCs we considered 752 scRNA-Seq profiles processed in two distinct runs using the Smart-seq2 protocol (J. Ding *et al*., 2019) **(Supplementary Table S3)**. Symphony based integration of the transcriptomes showed distinct localised cell populations, with ClearCell/Polaris enriched cells having no visible WBC contaminants. A fraction of CTCs from the Elbright dataset overlapped with WBCs **(Figure 5A).** Seurat identified a number of clusters with CTCs alone. However, it also returned a number of clusters with a fair bit of mixing of the two kinds **(Figure 5B)**. Clustering using DDLK grouped the CTCs into three clusters whereas the WBCs clumped into one large cluster **(Figure 5C)**. Notably, we found ClearCell FX and Polaris selected CTCs clustered with one of the ER+ subgroups sourced from the study by Ebright and colleagues (Ebright *et al*., 2020). Seurat and unCTC detected clusters and associated WBC-CTC distribution are depicted in **Figure 5D, E** and **Figure 5F, G** respectively. Out of the 72 CTCs that finally qualified the filtering criteria (obtained from ClearCell FX and Polaris workflow), ER+ cells were most prevalent (54 out of 72). Among the rest there were 7 and 11 cells of the HER2+ and triple negative categories respectively. One possible reason for not detecting HER2+ and triple negative CTCs as separate categories is their inadequate numbers, which makes it difficult for unCTC to retain relevant genes/pathways through several upstream filtering steps such as gene filtering and pathway selection.

**Figure 5.**
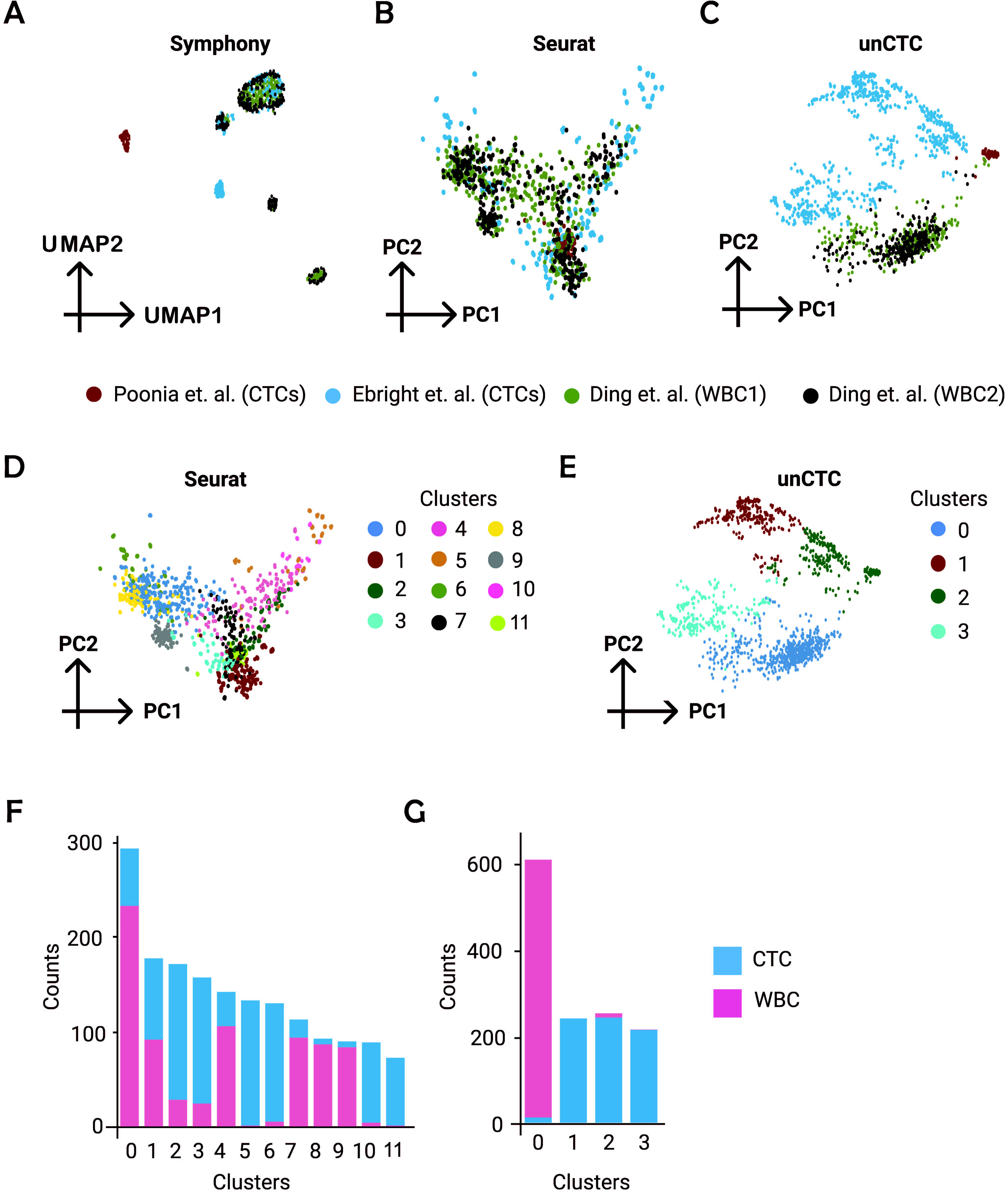
Clustering of CTCs obtained from ClearCell^®^ FX - Polaris^TM^ system. (A-C) Visualisation of ClearCell^®^ FX - Polaris^TM^ in presence of independent breast CTC and WBC scRNA-Seq profiles, using Symphony, Seurat and unCTC respectively. (D,E) Seurat and unCTC cluster identities. Notably, Symphony does not cluster single cells. unCTC accurately segregates CTCs and WBCs. CTCs obtained from ClearCell^®^ FX - Polaris^TM^ system co-cluster with breast CTCs from Ebright data (Ebright *et al*., 2020) . (F,G) CTC-WBC distribution across clusters detected by Seurat and unCTC respectively.

### Marker dependent characterisation of CTC clusters

Cell-lineages are best understood through enrichment of well-characterized lineage markers. Two approaches can be adopted for this. First, investigating differential expression for single markers, and second, for marker panels. For the second study (comprising ClearCell FX and Polaris) we analyzed lineage identities for clusters identified by DDLK **(Figure 6A)**. Multiple well-known immune-cell markers were spotted among the top 200 differentially upregulated genes **(Supplementary Table S5)** among cells in cluster 0 that predominantly contains WBCs from the Ding et al. dataset. These are *NKG7, PTPRC, PTPRCAP, IL32, CD74,* and *CD48*. Remaining clusters (clusters 1, 2, 3) comprise mostly CTCs (from Poonia and Ebright et al. datasets).

**Figure 6.**
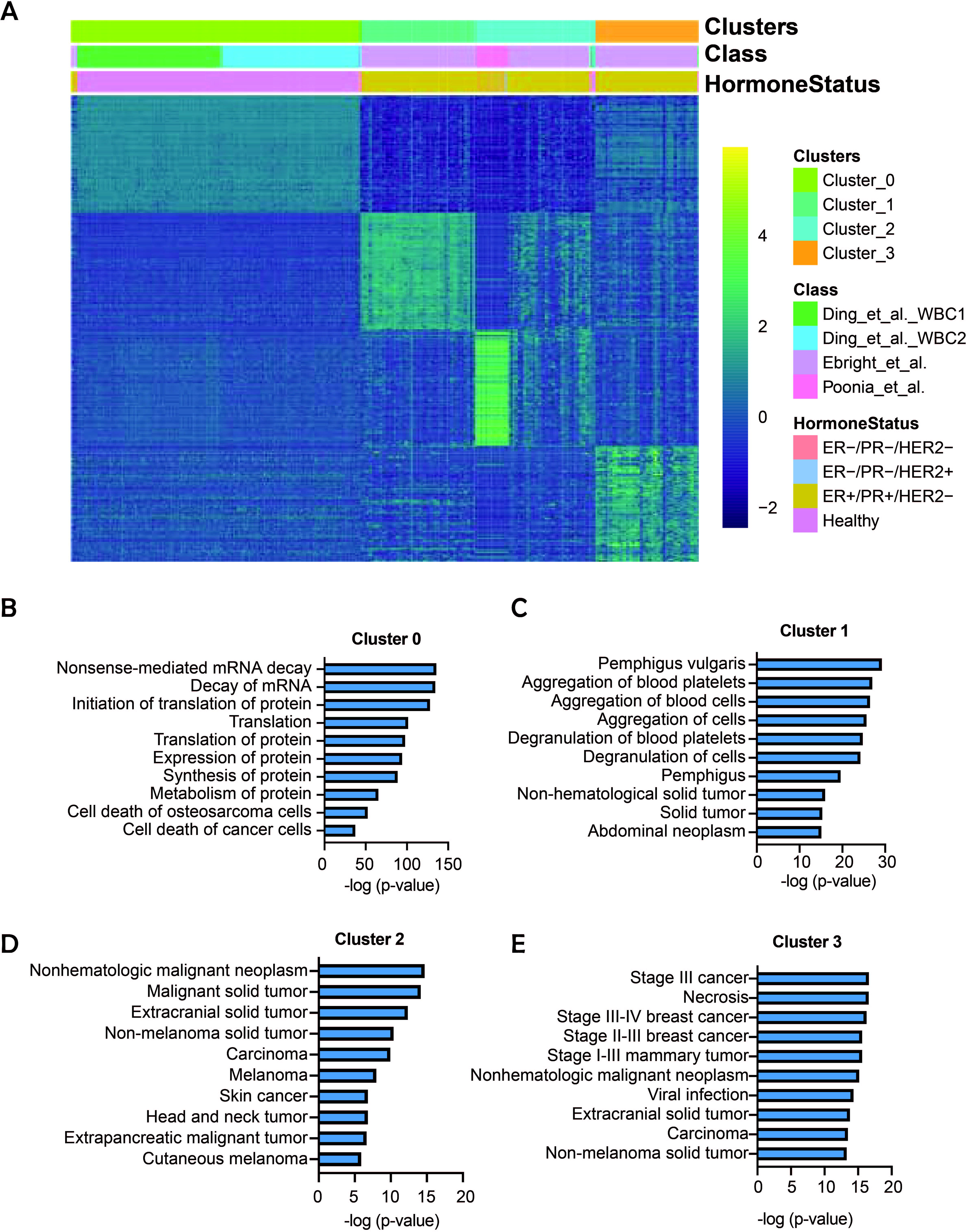
Functional annotations of the highly enriched circulating tumor cell-associated genes identified using unCTC. (A) Heatmap depicting the expression of top 200 differentially elevated genes across four clusters detected by unCTC. Color bars indicate cluster identity, source data information as well as molecular subcategories. (B-E) Bar plots depicting gene set enrichment using disease and functional annotation modules of IPA. The lists of cluster specific differentially elevated genes can be found in **Supplementary Table S5**.

Cluster 1 among these are found to have elevated expression levels of integrins (*ITGA2B* and *ITGB5*). Integrins are principal adhesion molecules and play a central role in platelet function and hemostasis. Recent studies have postulated CTC-platelet interaction based on RNA extracts of single and clustered CTCs (Szczerba *et al*., 2019; Aceto, 2020). CTCs constantly interact with factors in the blood such as platelets, circulating nucleic acids, and extracellular vesicles, which influence their molecular profiles (Ward *et al*., 2021). Interestingly, we observed elevated expression levels of platelet degranulation markers *CLU* and *SPARC*, which are known for regulating *PF4* (Beck *et al*., 2019), a critical endocrine factor previously described to be associated with worse outcome in patients with lung cancer (Pucci *et al*., 2016). *PF4* was also found to have elevated expression in cluster 1 specific cells. Cluster 1 specific CTCs showed elevated expression of numerous oncogenes with well-known roles in breast cancer progression. *CDKN1A (Koch et al., 2020)*, *TIMP1 (Abreu et al., 2020)*, and *PGRMC1 (Clark et al., 2016)* are notable among these. Cluster 2 that harbored the ClearCell/Polaris CTCs aside from CTCs from the Elbright dataset expressed. This cluster exhibited a number of breast cancer associated transcripts. Notable among these are — *IL10* that drives breast cancer progression and proliferation (Sheikhpour *et al*., 2018); *BRIP1,* whose genotypic alteration increases breast cancer risk and elevated expression features invasive nature of the primary disease (Eelen *et al*., 2008); IDO1 a key immune checkpoint protein (Dill *et al*., 2018), and *OCT4/POU5F1,* a cancer stem cell marker (Jin *et al*., 2019). Cluster 3 can be best characterised by elevation of canonical epithelial markers such as *EPCAM, KRT18,* and *KRT19.* Tumor suppressor *SOD1 (Liu et al., 2020)* was found to be upregulated in these cells.

Functional analysis of the cluster specific upregulated genes were performed using IPA (QIAGEN Inc., https://www.qiagenbioinformatics.com/products/ingenuitypathway-analysis) (Krämer *et al*., 2014). Cluster 0 (WBC cluster) specific upregulated genes were largely unrelated to cancer, whereas the remaining clusters (CTC specific) exhibited enrichment of cancer associated pathways aligned with our above analysis of cluster marker genes **(Figure 6B-E)**. We also visualized select relevant pathways that had differentially elevated enrichment in specific clusters **(Supplementary Figure S1)**. Cluster-specific differential enrichment of pathways was largely concordant with the insights gained based on analysis of differential expressed genes **(Supplementary Table S6)**.

The Poonia et al. dataset (ClearCell FX and Polaris) comes from ER-/PR-/HER2-, ER-/PR-/HER2+, and ER+/PR+/HER2-breast cancer patients. For each subtype, we identified the top 10 upregulated genes with a significant *P*-value and a Log Fold Change (LFC) > 2 **(Supplementary Figure S2, Supplementary Table S7)**. ER+/ PR+/ HER2− subtype showed elevated expression of *ZEB2*. Notably *ZEB2* has known roles in promoting metastasis and cell motility in estrogen receptor positive breast cancer (Burks et al. 2014). Another gene *ATF3* is known to be involved in promoting resistance to endocrine therapy in estrogen positive breast cancer (Borgoni et al. 2020). *EIF3C,* which is known to promote proliferation (Zhao *et al*., 2017), showed elevated expression levels in CTCs originating from the TNBC subtype.

We performed cluster characterization based on single markers and marker panels. It is well-known that due to the loss of epithelial property, only a tiny percentage of CTCs would display conventional epithelial markers (Iyer *et al*., 2020). Because of EMT in CTCs and high dropout rates in scRNA-Seq data, single markers may not show substantial differential expression; consequently, a combinatorial approach may be more beneficial than tracking differential expression of individual genes. We curated markers from literature, and these markers are highly expressed in immune cells and breast epithelia **(Supplementary Table S4)**. Stouffer’s method (Stouffer *et al*., 1949) was used for combinatorial scoring of marker enrichment at a single cell level. Scores obtained separated the WBC and CTC populations as expected **(Figure 7A)**. We then tracked differential expressions of single markers. Cluster 0 specific cells were found to express high levels of leukocyte markers such as *PTPRC* and *NKG7*. *EPCAM* and *KRT18* exhibited relatively higher expression levels in cells specific to clusters 1, 2, and 3 **(Figure 7B-E)**.

**Figure 7.**
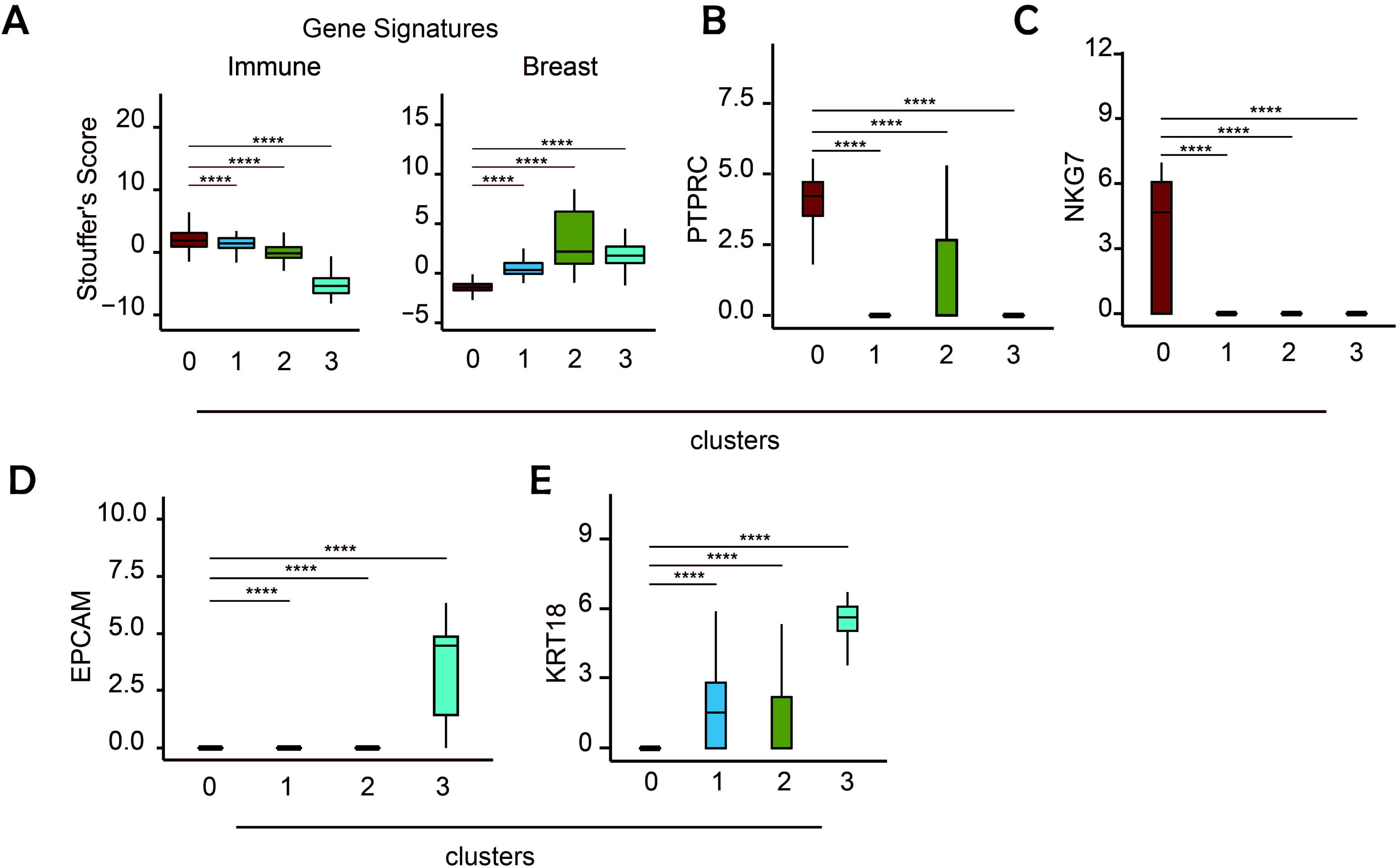
Lineage identity analysis using known markers and gene-sets. (A) Box plot depicting distribution of Stouffer’s scores (Stouffer *et al*., 1949) associated with genesets, specific to immune cells and breast epithelia **(Supplementary Table S4)**. Cluster 0 shows enrichment of immune cell specific markers whereas the remaining clusters show enrichment of markers specific to breast epithelia. (B, C) Box plots depicting differential enrichment of select immune cell markers i.e. PTPRC and NKG7 respectively. (D, E) Box plots depicting differential enrichment of select epithelial markers i.e. EPCAM and KRT18 respectively.

### Expression-based copy number variation inference

Duplications and deletions that result in the addition or loss of significant chromosomal regions are referred to as copy number variations (CNV). As proven by the Cancer Genome Atlas (Weinstein *et al*., 2013) and the International Cancer Genome Consortium (Mafficini and Scarpa, 2018), somatic CNVs, also known as copy number aberrations (CNAs), are prevalent in cancer. These CNAs are strongly linked to the onset, development, and metastasis of cancer (Sudmant *et al*., 2015; Jiang *et al*., 2016; Urrutia *et al*., 2018). With the growing popularity of single-cell sequencing of tumor microenvironments, expression based CNV inference has become critical in zeroing in on malignant cells in a marker-independent manner (Tickle *et al*., 2019). The same strategy can be useful to characterize CTCs.

We subjected the CTC and WBC clusters, including potential CTCs captured by ClearCell FX and Polaris workflow to inferCNV, a popular tool for CNV inference from single-cell expression data (Tickle *et al*., 2019). Our marker based analyses already highlighted cluster 0 as the one containing WBCs. We used transcriptomes from this cluster as reference for subtraction of the copy number signals from the remaining CTC rich clusters (clusters 1, 2, and 3) **(Figure 8)**. Since this method is confounded by clusters, we performed a separate inferCNV analysis considering ClearCell FX and Polaris selected CTCs separately from the rest of the CTCs **(Supplementary Figure S3)**. We spotted multiple locations of apparent copy number gains and losses in poonia et al. dataset. Past studies demonstrated that in breast cancer, typical chromosomal gains are on arms 1q32 and 1q42–q44 and chromosomal loss is rarely noticeable on the q arm (Orsetti *et al*., 2006). Chromosome 1q harbors both tumor suppressor genes and oncogenes and are linked with breast carcinogenesis. Two altered regions were identified in chromosome 1q: smallest commonly deleted and overrepresented regions at 1q21-31 and 1q41-q44 respectively (Bièche, Champème and Lidereau, 1995; Lobo, 2008); (Privitera, Barresi and Condorelli, 2021). Notably, we found deletions for tumor suppressor genes — Retinoid acid receptor beta2 (3p24), thyroid hormone receptor beta1 (3p24.3) and Ras association domain family 1A (3p21.3) genes in our in-house data (Ingvarsson, 2001); (Senchenko *et al*., 2004); (Ling *et al*., 2015). Previous studies have reported presence of CTC-like genomic gains in chromosome 19 at low frequency in primary breast cancer. Furthermore, several studies have suggested that gains in chromosome 19 may have a role in aggressive forms of breast cancer (Turner *et al*., 2010; Natrajan *et al*., 2012). 19q13 region is associated with copy number gains of signatures involved in dormancy and tumor aggressiveness in CTCs. Some of the genes present in these signatures are involved in promoting EMT, invasion, metastasis are *CEBPA* (19q13.11)*, FXYD5* (19q13.12)*, PAK4* (19q13.2)*, AKT2* (19q13.2) *(Kanwar et al., 2015).* We have found concordant indications from inferCNV-based analysis of the ClearCell FX and Polaris selected CTCs **(Supplementary Figure S3)**. A summary of each possible event and the CNV state (1 = 2 copy loss, 2 = 1 copy loss, 3 = neutral, 4 = 1 copy gain, 5 = 2 copy gain, 6 = 3+ copy gain) is given in the **Supplementary Table S8**, with source data identities.

**Figure 8.**
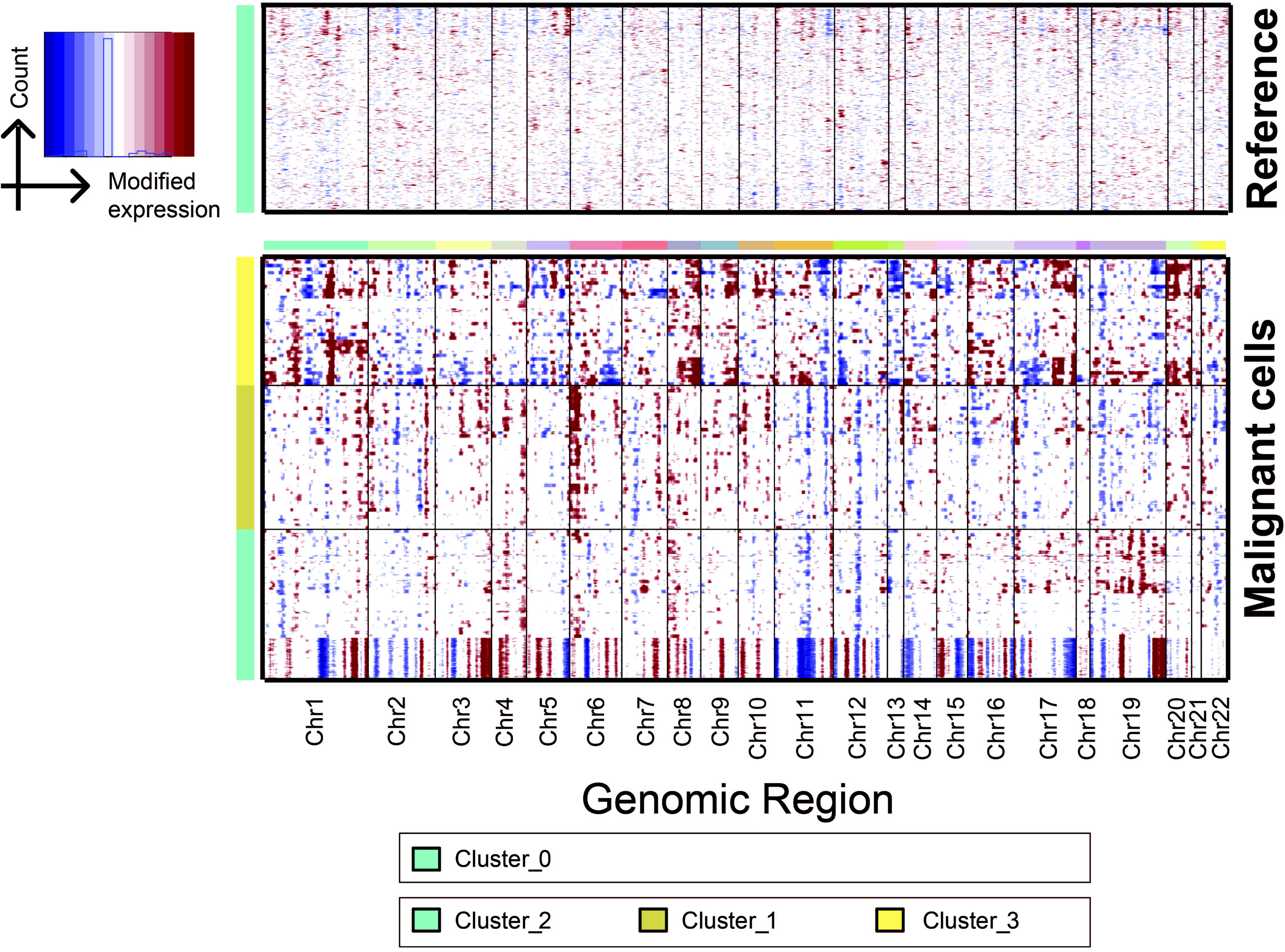
Expression based inference of CNV landscape across malignant cell clusters. Heatmap obtained from inferCNV tool (Tickle *et al*., 2019) depicting putative CNV landscape across malignant cell clusters, while considering cluster 0 (WBCs) as reference.

## Materials and Methods

### Description of datasets

We compiled two scRNA-Seq datasets for comprehensive evaluation of unCTC. In the first study, we used seven distinct single-cell RNA-Seq (scRNA-Seq) datasets of circulating tumor cells (CTCs) and white blood cells (WBCs) (Aceto *et al*., 2014; Ting *et al*., 2014; Yu *et al*., 2014; Sarioglu *et al*., 2015; Jordan *et al*., 2016; Velten *et al*., 2017; Zheng *et al*., 2017). Six of the seven datasets yielded 141 single CTCs. Two of the studies offered a total of 1037 WBCs. Notably, one of these datasets (accession number: GSE67939), contains both blood and CTC transcriptomes (Sarioglu *et al*., 2015). Another dataset (accession number: GSE74639) contains ten single CTCs and six single primary tumor cells (Zheng *et al*., 2017). The CTC data entails three cancer types: breast, lung, and pancreatic **(Supplementary Table S3)**. This dataset was used to validate the unCTC potential for integrative analysis and clustering.

In the second study, we utilized three publicly accessible scRNA-Seq datasets — (a) 81 potential CTCs enriched by the ClearCell FX and Polaris workflow. The CTCs were six patients with breast cancer of three subtypes—ER-/PR-/HER2-, ER+/PR+/HER2-, ER-/PR-/HER2+. This data is referred to as Poonia et al. dataset. (b) As control we also considered 824 ER+/PR+ single CTCs isolated directly from whole blood specimens of cancer patients using the CTC-iChip microfluidic system (Ebright et al. dataset) (Ebright *et al*., 2020). (c) A total of 752 WBC expression profiles (processed in two different runs). This dataset is referred to as Ding et al. dataset (J. Ding *et al*., 2019) **(Supplementary Table S3)**.

### Sample collection

In total 81 CTCs were collected from blood specimens of 6 breast cancer patients with distinct molecular subtypes. Out of these, 11 CTCs were obtained from one patient with TNBC, 57 CTCs from three patients with ER+/PR+/HER2-breast cancer and 13 CTCs from three patients with ER-/PR-/HER2+ subtype **(Supplementary Table S1)**. All blood samples were collected from breast cancer patients at National Cancer Centre Singapore with informed consent of all human participants and in accordance with institutional review board (IRB) guidelines (CIRB no. 2014/119/B). The SingHealth Centralised Institutional Review Board examined and approved the clinical sample collection protocols. The immunohistochemical (IHC) testing of estrogen receptor (*ER*), progesterone receptor (*PR*) and human epidermal growth factor receptor 2 (*HER2*) status was conducted based on the the latest guidelines of the American Society of Clinical Oncology and the College of American Pathologists.

### CTC enrichment

To perform CTC enrichment, 9 mL of blood samples were collected in K3 EDTA blood collection tubes (Greiner Bio-One, 455036). For each run, 6–8.5 mL of whole blood was processed. The red blood cells were first removed with addition of red blood cell (RBC) lysis buffer (G-Biosciences®, St. Louis, MO, USA) followed by incubation of 10 min at room temperature. After centrifugation, the lysed RBCs in the supernatant were removed. The nucleated cell pellet was suspended in a ClearCell resuspension buffer before CTC enrichment on the ClearCell FX system (Biolidics Limited)(Lee, Guan and Bhagat, 2018) as per the manufacturer’s instructions.

### Immunofluorescence suspension staining

CTC-rich blood samples were centrifuged at 300 *g* for 10 min and concentrated to 70 μL. The cell staining was performed with the addition of the following markers and antibodies for one hour: CellTracker™ Orange (CTO) (Thermo Fisher, C34551), Calcein AM (Thermo Fisher, L3224), *CD45* antibody conjugated with Alexa 647 (BioLegend®, 304020), and *CD31* conjugated with Alexa 647 (BioLegend, 303111). To improve the viability and RNA quality of the cells, 15 μL of RPMI with 10% FBS (Gibco) and 3 μL of RNase inhibitor (Thermo Fisher, N8080119) were also added. After incubation, 13 mL of PBS was added to dilute the staining reagents. The sample was spun down at 300 *g* for 10 min and concentrated to 45 μL. In order to achieve optimal buoyancy in an integrated fluidic circuit (IFC), 45 μL of CTCs was mixed with a 30 μL Cell Suspension Reagent (Fluidigm, 101-0434) to achieve 75 μL of cell mix.

### Integrated fluidic circuit (IFC) operation

The Polaris IFC was first primed using the Fluidigm Polaris system to fill the control lines on the fluidic circuit, load cell capture beads, and block the inside of polydimethylsiloxane (PDMS) channels to avoid non-specific absorption/adsorption of proteins. Then to capture and maintain the single cells in the sites, capture sites (48 sites) were preloaded with beads that are coupled on IFC to build a tightly packed bead column during the IFC priming process. After the priming stage, the cell mix (cells with suspension reagent) was laden in three inlets (25 μL each of cell mix) on the Polaris IFC and single CTO+ & Calcein AM+ & CD45− & CD31− cells were selected to capture sites. Finally, single-cell processing was achieved through template-switching mRNA-Seq chemistry for full-length cDNA generation and pre-amplification on IFC.

### mRNA-Seq library preparation and sequencing

SMARTer^®^ Ultra^®^ Low RNA Kit for Illumina^®^ Sequencing (Clontech^®^, 634936) was employed to generate pre-amplified cDNA. The Polaris cell lysis mixture was used to lyse the selected and sequestered single cells. The 28 μL cell lysis mix is composed of 8.0 μL of Polaris Lysis Reagent (Fluidigm, 101-1637), 9.6 μL of Polaris Lysis Plus Reagent (Fluidigm, 101-1635), 9.0 μL of 3 SMART CDS Primer II A (12 M, Clontech, 634936), and 1.4 μL of Loading Reagent (20X, Fluidigm, 101-1004). The thermal profile for single-cell lysis is 37 °C for 5 min, 72 °C for 3 min, 25 °C for 1 min, and hold at 4 °C. The 48 μL preparation volume for reverse transcription (RT) contains 1X SMARTer Kit 5X First-Strand Buffer (5X; Clontech, 634936), 2.5-mM SMARTer Kit Dithiothreitol (100 mM; Clontech, 634936), 1 mM SMARTer Kit dNTP Mix (10 mM each; Clontech, 634936), 1.2 μM SMARTer Kit SMARTer II A Oligonucleotide (12 μM; Clontech, 634936), 1 U/μL SMARTer Kit RNase Inhibitor (40 U/μL; Clontech, 634936), 10 U/μL SMARTScribe™ Reverse Transcriptase (100 U/μL; Clontech, 634936), and 3.2 μL of Polaris RT Plus Reagent (Fluidigm, 101-1366). All the concentrations correspond to those found in the RT chambers inside the Polaris IFC. The thermal protocol for RT is 42 °C for 90 min (RT), 70 °C for 10 min (enzyme inactivation), and a final hold at 4 °C.

The 90 μL preparation volume for PCR contains 1X Advantage® 2 PCR Buffer [not short amplicon (SA)](10X, Clontech, 639206, Advantage 2 PCR Kit), 0.4-mM dNTP Mix (50X/10 mM, Clontech, 639206), 0.48-μM IS PCR Primer (12 μM, Clontech, 639206), 2X Advantage 2 Polymerase Mix (50X, Clontech, 639206), and 1X Loading Reagent (20X, Fluidigm, 101-1004). All the concentrations correspond to those found in the PCR chambers inside the Polaris IFC. The thermal protocol for preamplification consists of 95 °C for 1 min (enzyme activation), five cycles (95 °C for 20 s, 58 °C for 4 min, and 68 °C for 6 min), nine cycles (95 °C for 20 s, 64 °C for 30 s, and 68 °C for 6 min), seven cycles (95 °C for 30 s, 64 °C for 30 s, and 68 °C for 7 min), and final extension at 72 °C for 10 min. The preamplified cDNAs are harvested into 48 separate outlets on the Polaris IFC carrier. The cDNA reaction products were then converted into mRNA-Seq libraries using the Nextera^®^ XT DNA Sample Preparation Kit (Illumina, FC-131-1096 and FC-131-2001, FC-131-2002, FC-131-2003, and FC-131-2004) following the manufacturer’s instructions with minor modifications. Specifically, reactions were run at one-quarter of the recommended volume, the tagmentation step was extended to 10 min, and the extension time during the PCR step was increased from 30 to 60 s. After the PCR step, samples were pooled, cleaned twice with 0.9× Agencourt® AMPure® XP SPRI beads (Beckman Coulter), eluted in Tris + EDTA buffer and quantified using a high-sensitivity DNA chip (Agilent). The pooled library was sequenced on Illumina NextSeq® using reagent kit v3 (2 × 74 bp paired-end read).

### Preprocessing of scRNA-Seq datasets

The unCTC R package accepts scRNA-Seq data in two forms: transcripts per million (TPM) and raw count data. In the first study, we downloaded scRNA-Seq count data from all respective sources. The second study includes three different datasets, including our own. Ding et al. dataset (raw count) was downloaded from the Broad Institute’s single-cell gateway. Ebright et al. and Poonia et al. datasets were obtained by processing the associated FASTQ files. We used the FastQC tool to perform quality checks on both datasets for average percent GC content, mean quality score, and per-sequence quality score (Andrews, 2010). For alignment purposes of Ebright data set, we used the hg19 reference genome and hg19 GTF file from Ensembl (release 75) (Howe *et al*., 2021). To estimate the expression levels of genes, we used RNA-Seq by Expectation-Maximization. v.1.3.1 (RSEM) (Li and Dewey, 2011) with two scripts: *rsem-prepare-reference* and *rsem-calculate-expression*. Finally, length-normalized TPM datasets (reporting expression of 57773 transcripts) were obtained for both studies. **Supplementary Figure S4** shows the steps used in preprocessing of the single-cell RNA-Seq datasets. For alignment purposes of Poonia data set, An index for RNA-Seq by expectation maximization (RSEM) was generated based on the hg19

RefSeq transcriptome downloaded from the UCSC Genome Browser database. Read data were aligned directly to this index using RSEM/bowtie. Quantification of gene expression levels in counts for all genes in all samples was performed using RSEM v1.2.4. Genomic mappings were performed with TopHat 2 v2.0.13, and the resulting alignments were used to calculate genomic mapping percentages. Raw sequencing read data were aligned directly to the human rRNA sequences NR_003287.1 (28s), NR_003286.1 (18S) and NR_003285.2 (5.8S) using bowtie 2 v2.2.4.

### Data integration, filtration, and normalization

RSEM software returns both read count data and TPM data (Poonia et al. and Ebright et al.) From these datasets, we discarded cells with a total read-count less than 50000. As per this criterion, 9 CTCs were removed from the Poonia et al. scRNA-Seq dataset **(Supplementary Table S2)**. All 824 cells in the Ebright et al. dataset qualified this criterion. Poonia and Ebright et al. datasets were integrated with Ding et al. dataset containing WBC expression profiles and genes that are common across the datasets were retained. Further gene/cell filtering steps were implemented as follows. We eliminated cells with fewer than 1500 expressed genes (non-zero read count). On the other hand we considered genes with non-zero expression in at least five cells. Linnorm normalization technique method was used with default parameters for single-cell normalization and batch correction (Yip *et al*., 2017). Normalized expression values are log-transformed after the addition of 1 as pseudo count. Linnorm normalization and log transformation are applied to both count matrix and TPM matrix.

### Expression values to pathway enrichment scores

For computing gene-set enrichment scores we used the Gene Set Variation Analysis (GSVA) R software package (Hänzelmann, Castelo and Guinney, 2013). GSVA needs mainly two inputs: normalized and log-transformed expression matrix and gene sets. We used the C2 collection from MSigDB (Subramanian *et al*., 2005). This contains over 6000 literature-curated gene-sets. Before passing the geneset and expression matrix to the GSVA function, a filtering step is applied on genesets that removes genes that are not present in the normalized expression matrix. We set *min.sz* as 10, *max.sz* as 500, *max.diff* as FALSE. Since the calculations for each gene-set are independent of each other, we calculated enrichment scores in parallel. Here we used four parallel threads to speed up the computation (parallel.sz=4).

### DDLK clustering

Unsupervised clustering of GSVA enrichment scores was performed using K-means friendly deep dictionary learning (DDL). To specify the optimal value of clusters for K-means clustering, the elbow method is used.

The popular way to express K-means clustering is via the following formulation:

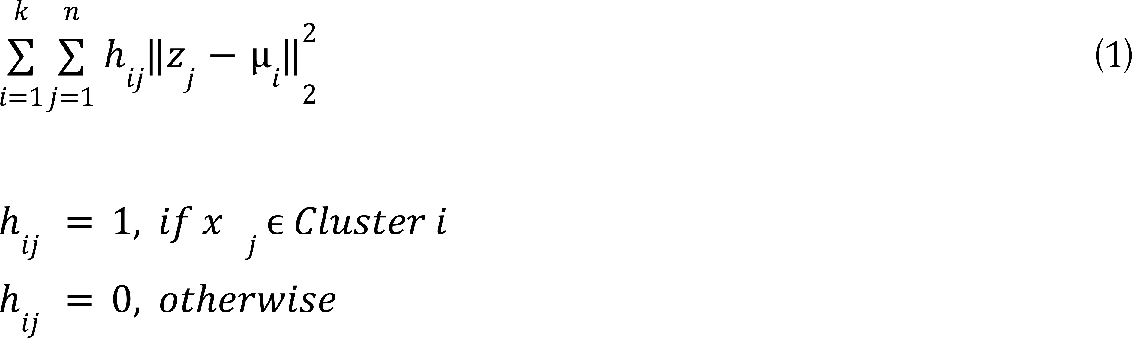

where z_j_ denotes the j^th^ sample and µ_i_ the i^th^ cluster.

An alternate formulation for K-means is via matrix factorization (Bauckhage, 2015).

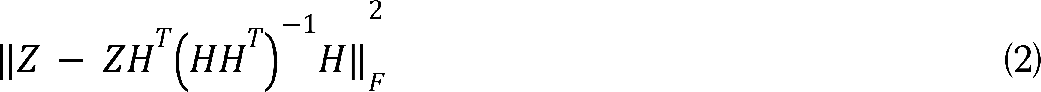

where Z is the data matrix formed by stacking z_j_’s as columns and H is the matrix of binary indicator variables h_ij_. We prefer expressing K-means as (2) in this work.

Since DDL is not a popular framework, we review it briefly. Dictionary learning (Tošić and Frossard, 2011) learns a basis (D) such that the data (X) can be generated / synthesized from the coefficients (Z).

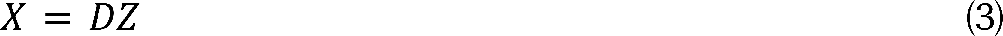

The term dictionary learning is relatively new. The same problem has been known as matrix factorization in the past. One can see that (3) is factoring the data matrix X into D and Z. In its most basic form, dictionary learning / matrix factorization is solved via the following –

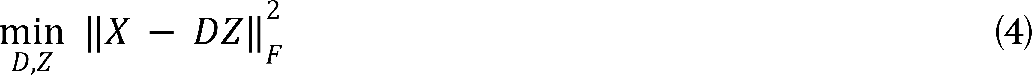

where 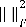 is the squared Frobenius norm defined as the sum of the squares of all the terms in the matrix.

In DDL, instead of learning one layer of the dictionary, multiple layers are learned instead. This is expressed as,

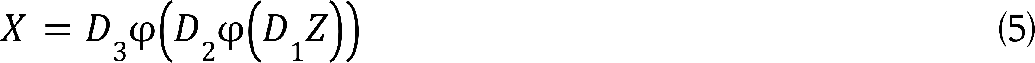

Here D_1_, D_2_, D_3_ are three layers of dictionaries and φ is the activation function between two layers. It is shown for three layers as an example, it can be more than three.

The solution to the unsupervised formulation is expressed as follows –

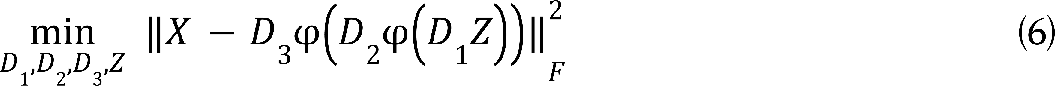

In [9], a greedy solution to (6) was proposed. This was not optimal in the sense that there was feedback from shallow to deeper layers but not vice versa. To overcome this the joint solution was proposed in (Singhal and Majumdar, 2018) based on the majorization minimization (MM) approach.

In this work, we will use the ReLU activation function for two reasons – 1. It is easier to incorporate as an optimization constraint, and 2. ReLU has been proven to have better function approximation capabilities. Therefore our basic framework for DDL (with ReLU) will be expressed as follows,

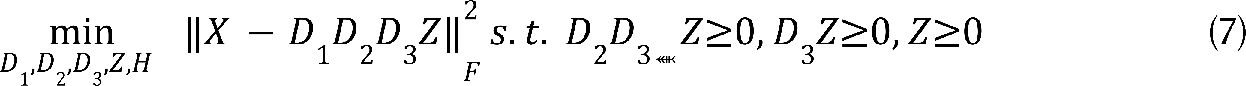

We propose to incorporate the K-means cost (2) into the DDL formulation (7). The basic idea is to use the features generated by DDL as inputs for clustering. However, instead of solving it piecemeal we jointly optimize the following cost function –

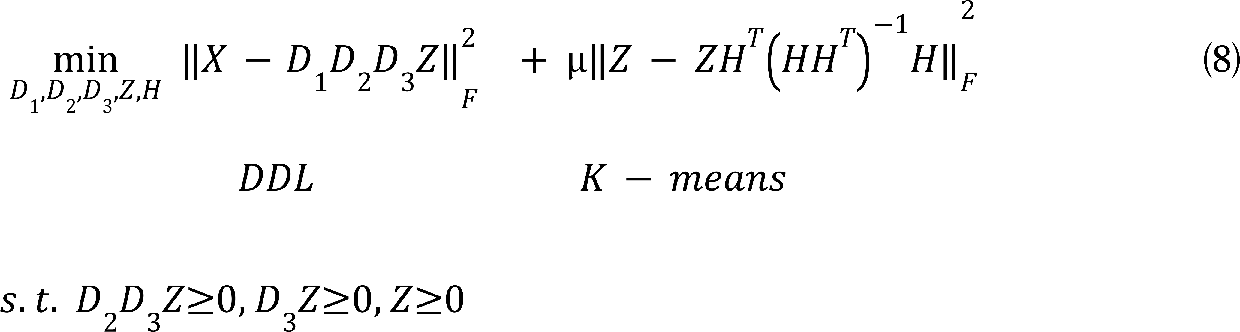

We solve (8) using alternating minimization. Initially, we ignore the non-negativity constraints in (8); later on, we will discuss how they can be handled. The updates for different variables are as follows,

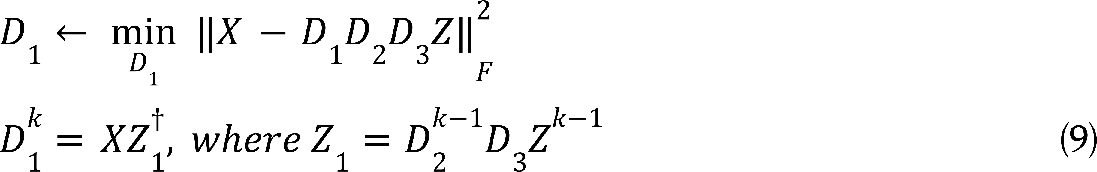

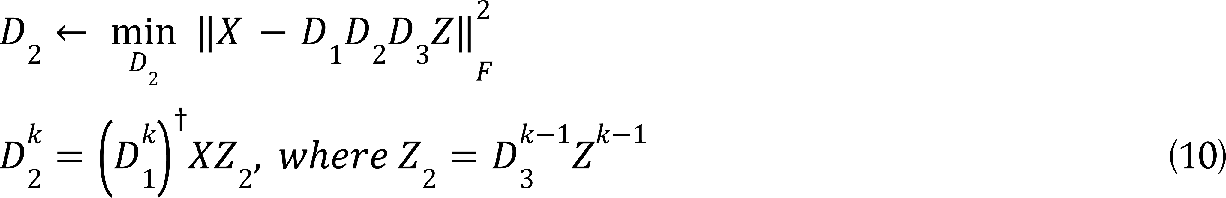

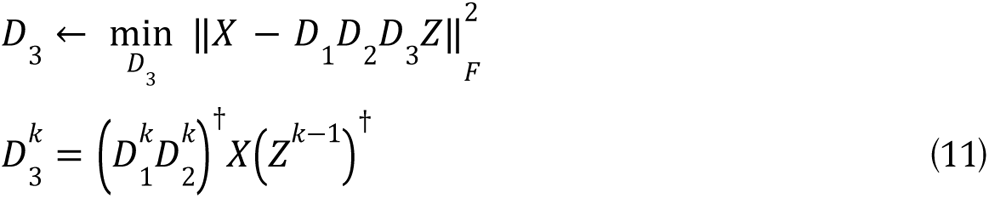

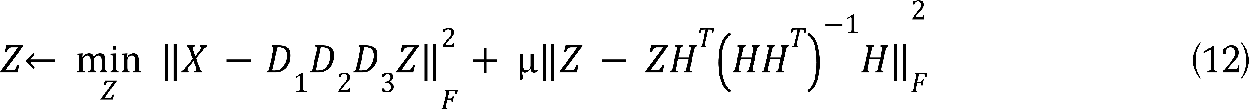

To solve Z, we need to take the gradient of the expression in (12) and equate it to zero. The derivation is given below.

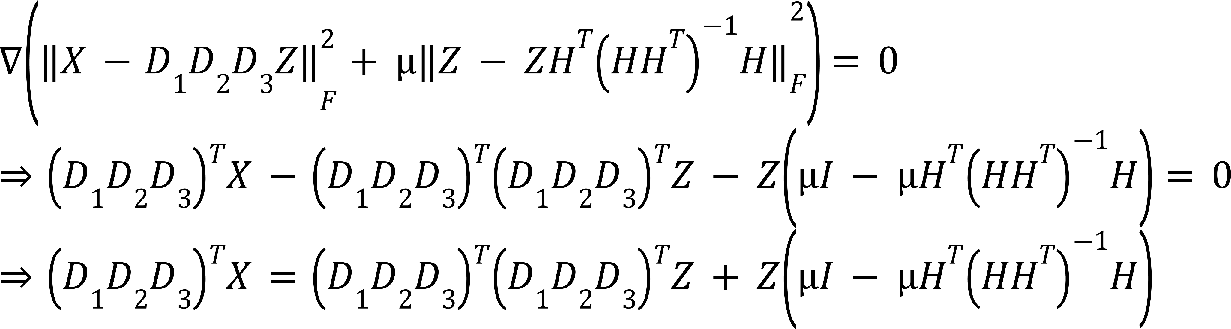

The last step of the derivation implies that Z is a solution to Sylvester’s equation of the form AX + XB = C. There are many efficient solvers for the same.

The final step is to update H. This is obtained by solving,

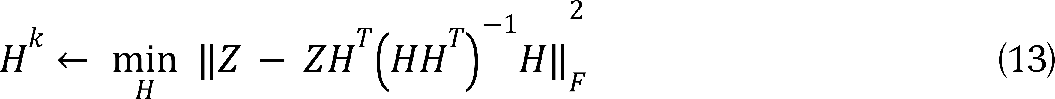

This is the K-means algorithm applied on Z.

In the derivation so far, we have not accounted for the ReLU non-negativity constraints. Ideally imposing the constraints would require solving them via forward-backward type splitting algorithms; such algorithms are iterative and hence would increase the run-time of the algorithm. We account for these constraints by simply putting the negative values in Z, Z_1_ and Z_2_ to zeroes in every iteration.

The algorithm is shown in a succinct fashion below. Once the convergence is reached, the clusters can be found from H. Since (8) is a non-convex function, we do not have any guarantees for convergence. We stop the iterations when the H does not change significantly in subsequent iterations.

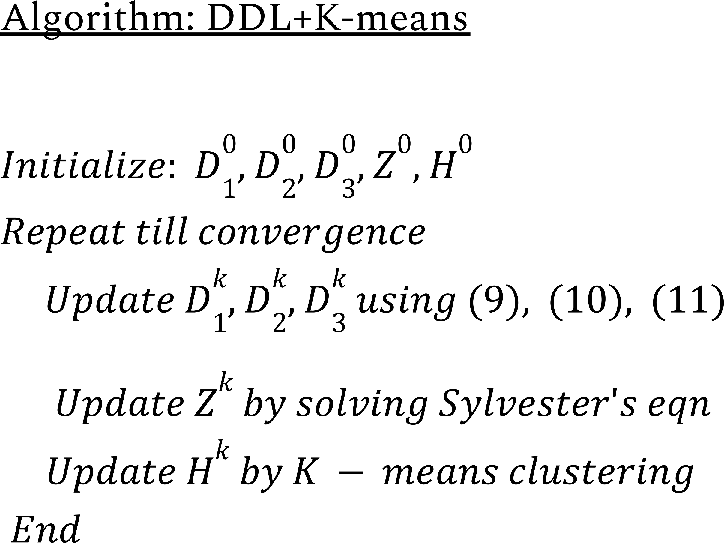

### DEGs identification

Differential genes between clusters obtained from DDLK clustering were calculated using the Limma (Ritchie *et al*., 2015) package with its voom (Law *et al*., 2014) method. We first used the normalized expression matrix to construct a DEGlist object. The DEGlist object was passed to the *calcNormFactors()* function of the edgeR R package (Robinson, McCarthy and Smyth, 2010), while setting *normalisation factor = 1* and *method = none,* followed by Voom transformation. We reported top 200 upregulated genes in each cluster that qualify adjusted p-value and log fold change (LFC) thresholds and then functional analysis of the cluster specific upregulated genes were performed using Ingenuity Pathway Analysis (IPA) (Krämer *et al*., 2014).

### Differential pathways

The R package Limma was used to obtain differential pathways between clusters identified by the DDLK clustering (Ritchie *et al*., 2015). The moderated t-statistic was used for differential pathway analysis. Pathways with positive log fold change and adjusted *P*-value < 0.05 were considered specifically enriched in a cell-group.

### Copy number variation analysis at chromosomal p or q arm

To infer copy number variation (CNV) landscape in single cancer cells, we used the inferCNV R package (Tickle *et al*., 2019). WBC cluster (identified through marker based analysis) was used as a healthy reference to estimate copy number aberrations in the CTC clusters. InferCNV plot shows significant CNVs between the control and test scRNA-Seq profiles. We used 1 as cutoff for the minimum average read counts per gene among reference cells, clustered according to the annotated cell types. To infer the location of a particular gain or loss on a chromosome at p or q-arm level, we used the cytoband information based on GRCh37 (Barrios and Prieto, 2017).

### Combinatorial evaluation of lineage markers

Stouffer’s method allows combining Z scores across multiple variables to arrive at a single score indicating enrichment of a certain property (or phenotype) (Gupta *et al*., 2020). We used this to measure enrichment of a spectrum of lineage indicating genes (breast epithelial and immune cell subtypes). This is particularly helpful for CTCs, since single markers may not exhibit adequate statistical significance for differential expression.

## Conclusion

Circulating tumor cells provide a window into their respective tumors of origin and cancer evolution. Most of the existing CTC enrichment methods are incomprehensive since they miss CTCs that do not express canonical epithelial markers. Single-cell expression studies allow inspection of molecular profiles of CTC rich cell populations obtained from an enrichment device. To date, there is no comprehensive computational resource that allows identification of diverse/unknown CTC phenotypes from scRNA-Seq data, comprising CTCs and WBCs. unCTC enables this by providing a number of unsupervised and semi-supervised means to interrogate CTC and WBC transcriptomes.

We demonstrated DDLK, a novel clustering approach that leverages pathway enrichment scores to yield robust grouping of single cells, even when the datasets are sourced from disparate studies. This is particularly helpful since typical single-cell studies feature multiple replicates. It should be noted that DDLK is meant to discover broad groups in an scRNA-Seq data. It is neither tested nor expected to aid discovery of heterogeneous subpopulations. For that one can refer to our previous works describing the dropClust software suite (Sinha *et al*., 2018, 2019). With the help of unCTC, we could spot CTCs, with unknown phenotypes. Expression based CNV inference corroborated our findings with precise genomic locations indicating amplification/deletion that are previously known in breast cancers.

CTCs are crucial biomarkers to monitor cancer progression and treatment response. Given the increasing throughput and sharply dropping price associated with single cell expression profiling, we predict unCTC will play an important role in constructing cancer specific molecular atlas of CTCs.

## Data availability

All raw and processed sequencing data generated in this study have been submitted to the NCBI Gene Expression Omnibus (GEO; https://www.ncbi.nlm.nih.gov/geo/) under accession number GSE186288 (https://www.ncbi.nlm.nih.gov/geo/query/acc.cgi?acc=GSE186288), reviewers’ token: sfobcuwizhsbrcl).

Download package and code from: https://github.com/SaritaPoonia/unCTC

## Author contribution

DS conceived the study with NR and AM. SP carried out all computational analyses with assistance from SC, NB and PR. AM, along with DS supervised machine learning method development. SP implemented the method with help from AG. NR, JW, and AAB conceived the integration of ClearCell^®^ FX and Polaris^TM^. NR and YFL developed the marker-free workflow. YSY provided the patient samples. YFL tested patient samples. GA assisted in interpreting results along with JT and AM. GA ideated and improved the scientific illustrations. All authors contributed in writing and proof-reading the manuscript.

## Supporting information

Supplementary Table S3: Description of datasets.

Supplementary Table S4: Highly expressed markers in immune cells and epithelial cells.

Supplementary Table S5: Cluster wise differentially upregulated genes.

Supplementary Table S6: Clusterwise differentially enriched pathways

Supplementary Table S7: Differential genes across CTCs of three different subtypes.

Supplementary Table S8: CNV events in CTCs.

---

**Figure S1.**
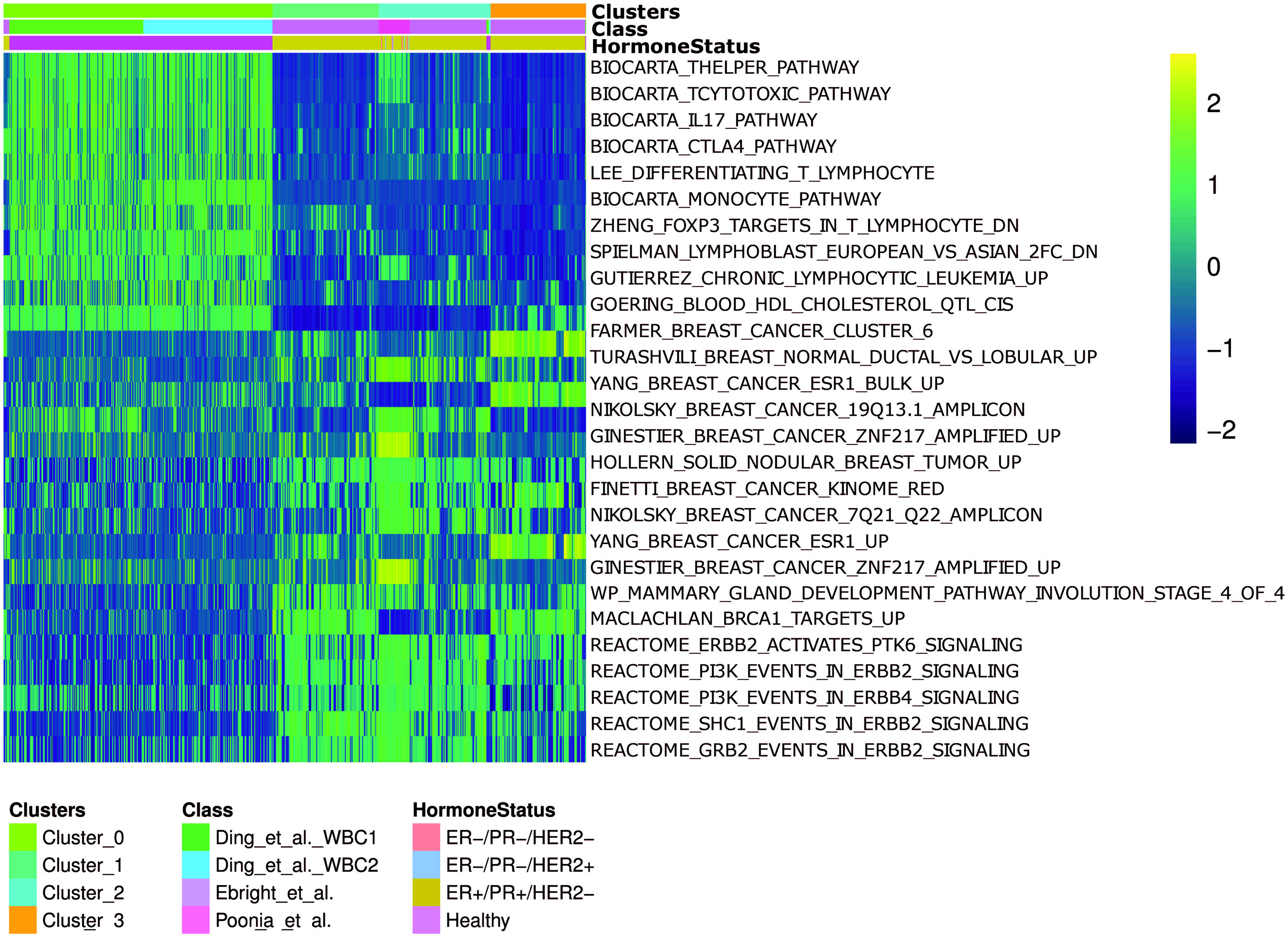
Heatmap of differential pathway enrichment scores across four clusters detected by unCTC. Color bars indicate cluster identity, source data information as well as molecular subcategories.

**Figure S2.**
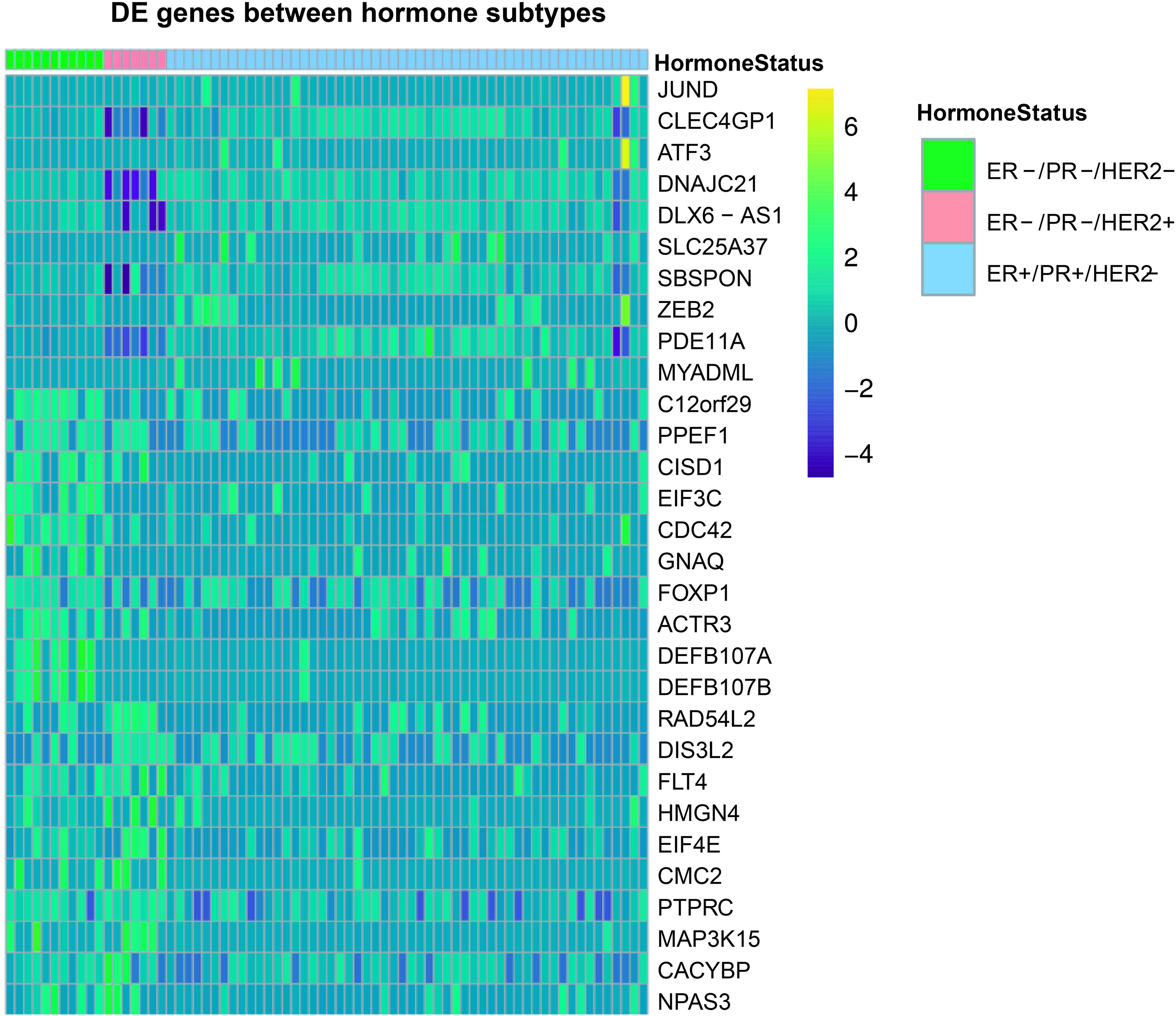
Heatmap depicting differential gene expression across ClearCell® FX and Polaris^TM^ selected CTCs across three molecular subtypes.

**Figure S3.**
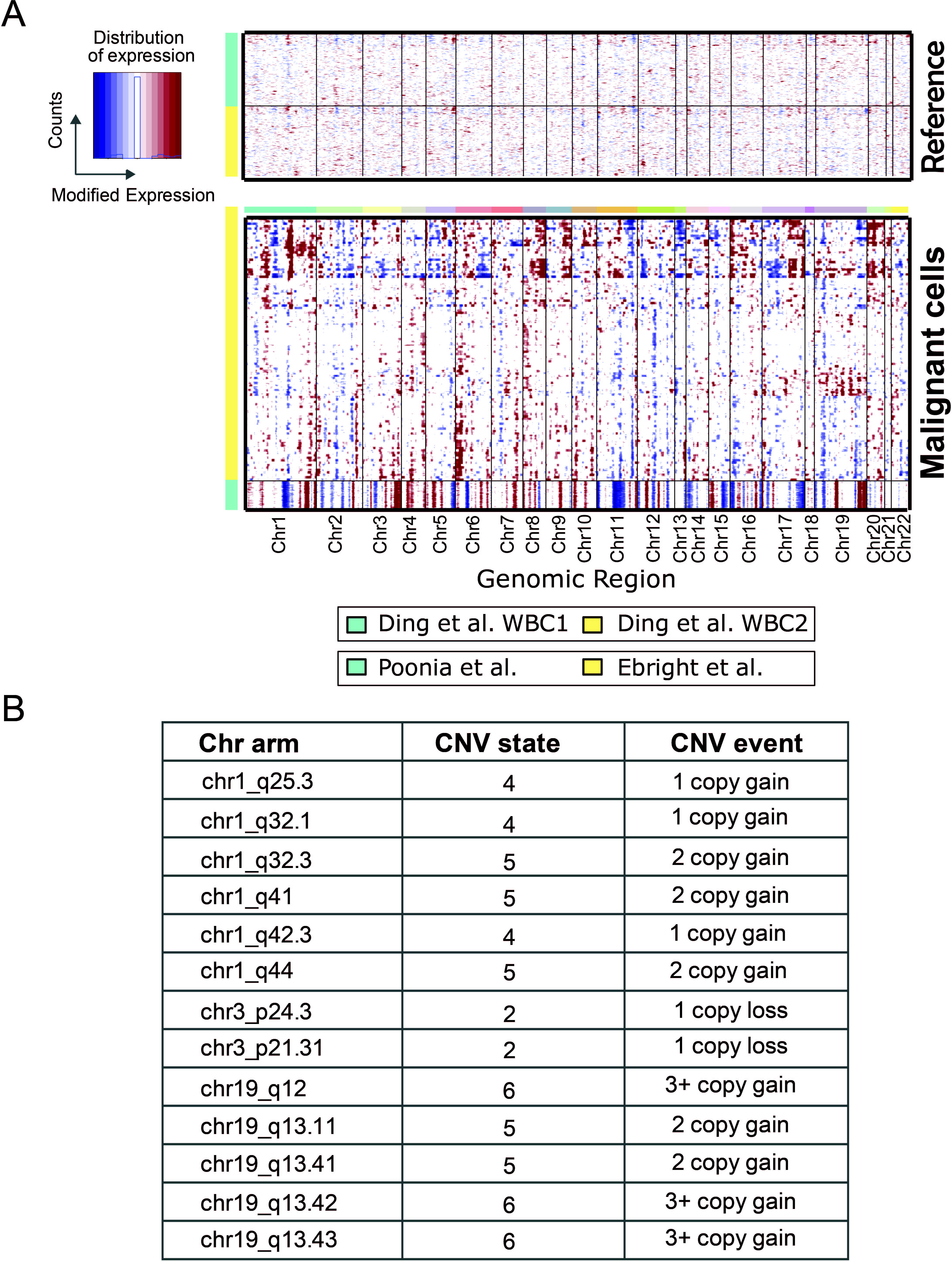
(A) Heatmap obtained from inferCNV tool (Tickle *et al*., 2019) depicting putative CNV landscape across CTCs obtained from ClearCell® FX and Polaris^TM^ system (referred to as Poonia dataset) as well as CTCs from the Ebright dataset. Cluster 0 (WBCs) is used as reference. (B) Table depicting some breast cancer relevant CNVs with their copy number states and precise genomic locations.

**Figure S4.**
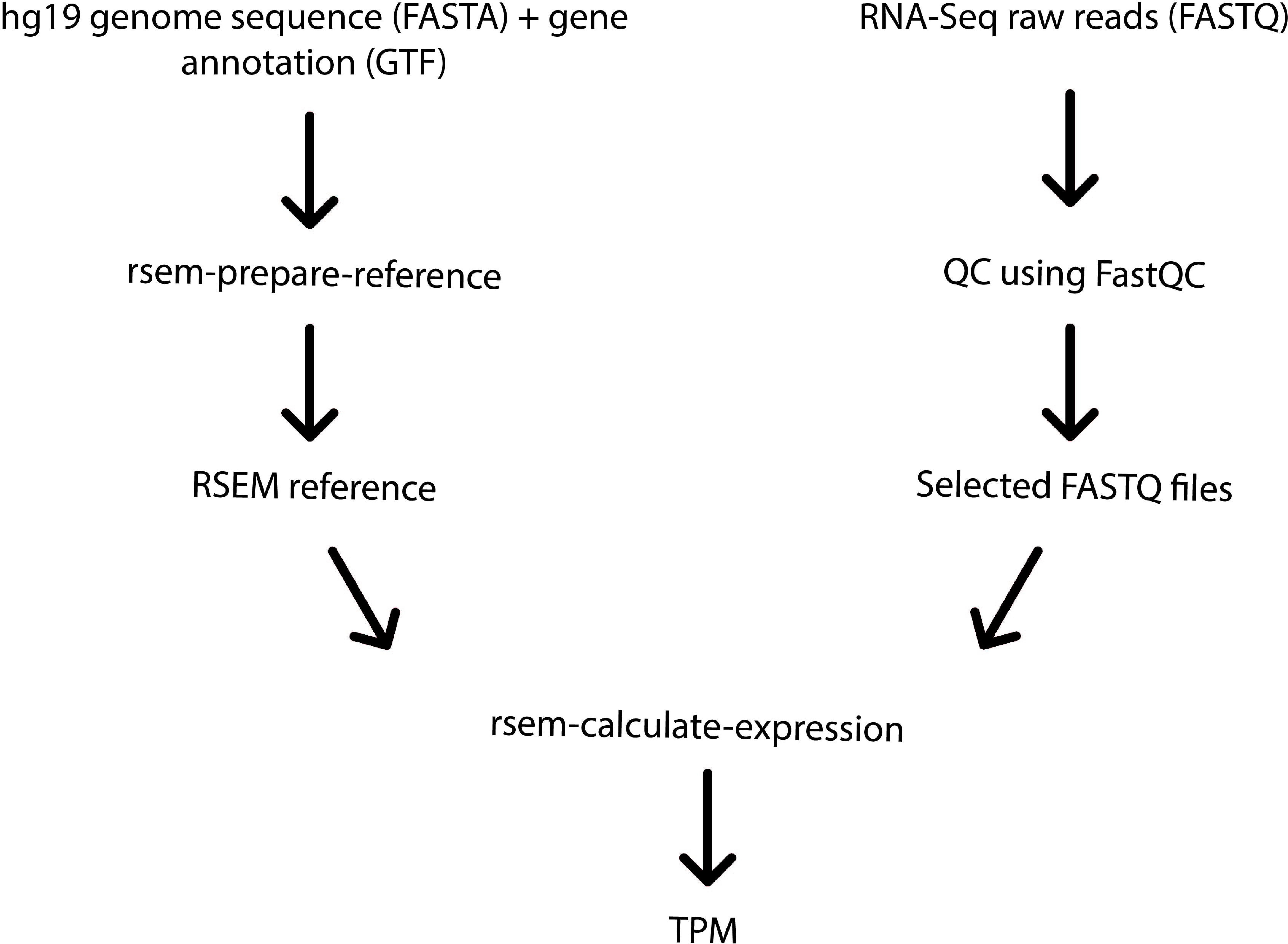
Computational workflow depicting key steps involved in the generation of TPM matrix using the raw RNA-Seq FASTQ files.

## References

Abreu, M. et al. (2020) ‘Circulating Tumor Cells Characterization Revealed TIMP1 as a Potential Therapeutic Target in Ovarian Cancer’, Cells , 9(5). doi: 10.3390/cells9051218.

Aceto, N. et al. (2014) ‘Circulating tumor cell clusters are oligoclonal precursors of breast cancer metastasis’, Cell, 158(5), pp. 1110–1122.

Aceto, N. (2020) ‘Bring along your friends: Homotypic and heterotypic circulating tumor cell clustering to accelerate metastasis’, Biomedical journal, 43(1), pp. 18–23.

Alix-Panabières, C. and Pantel, K. (2013) ‘Circulating tumor cells: liquid biopsy of cancer’, Clinical chemistry, 59(1), pp. 110–118.

Andrews, S. (2010) ‘Babraham bioinformatics-FastQC a quality control tool for high throughput sequence data’, URL: https://www.bioinformatics.babraham.ac.uk/projects/fastqc.

Barrios, D. and Prieto, C. (2017) ‘D3GB: An Interactive Genome Browser for R, Python, and WordPress’, Journal of computational biology: a journal of computational molecular cell biology, 24(5), pp. 447–449.

Bauckhage, C. (2015) ‘k-Means Clustering Is Matrix Factorization’, arXiv [stat.ML]. Available at: http://arxiv.org/abs/1512.07548.

Beck, T. N. et al. (2019) ‘Circulating tumor cell and cell-free RNA capture and expression analysis identify platelet-associated genes in metastatic lung cancer’, BMC cancer, 19(1), p. 603.

Bidard, F.-C., Proudhon, C. and Pierga, J.-Y. (2016) ‘Circulating tumor cells in breast cancer’, Molecular oncology, 10(3), pp. 418–430.

Bièche, I., Champème, M. H. and Lidereau, R. (1995) ‘Loss and gain of distinct regions of chromosome 1q in primary breast cancer’, Clinical cancer research: an official journal of the American Association for Cancer Research, 1(1), pp. 123–127.

Bittner, K. R., Jiménez, J. M. and Peyton, S. R. (2020) ‘Vascularized Biomaterials to Study Cancer Metastasis’, Advanced healthcare materials, 9(8), p. e1901459.

Bork, U. et al. (2015) ‘Circulating tumour cells and outcome in non-metastatic colorectal cancer: a prospective study’, British journal of cancer, 112(8), pp. 1306–1313.

Bulfoni, M. et al. (2016) ‘Dissecting the Heterogeneity of Circulating Tumor Cells in Metastatic Breast Cancer: Going Far Beyond the Needle in the Haystack’, International journal of molecular sciences, 17(10). doi: 10.3390/ijms17101775.

Butler, A. et al. (2018) ‘Integrating single-cell transcriptomic data across different conditions, technologies, and species’, Nature biotechnology, 36(5), pp. 411–420.

Büttner, M. et al. (2017) ‘Assessment of batch-correction methods for scRNA-seq data with a new test metric’, BioRxiv. Available at: https://www.biorxiv.org/content/10.1101/200345v1.abstract.

Chawla, S. et al. (2021) ‘UniPath: a uniform approach for pathway and gene-set based analysis of heterogeneity in single-cell epigenome and transcriptome profiles’, Nucleic acids research, 49(3), p. 1801.

Cheng, Y.-H. et al. (2019) ‘Hydro-Seq enables contamination-free high-throughput single-cell RNA-sequencing for circulating tumor cells’, Nature communications, 10(1), p. 2163.

Chen, L. et al. (2020) ‘Integrating Deep Supervised, Self-Supervised and Unsupervised Learning for Single-Cell RNA-seq Clustering and Annotation’, Genes, 11(7). doi: 10.3390/genes11070792.

Chiu, T.-K., et al. (2016) ‘Application of optically-induced-dielectrophoresis in microfluidic system for purification of circulating tumour cells for gene expression analysis-Cancer cell line model’, Scientific reports, 6, p. 32851.

Clark, N. C. et al. (2016) ‘Progesterone receptor membrane component 1 promotes survival of human breast cancer cells and the growth of xenograft tumors’, Cancer biology & therapy, 17(3), pp. 262–271.

Couturier, C. P. et al. (2020) ‘Single-cell RNA-seq reveals that glioblastoma recapitulates a normal neurodevelopmental hierarchy’, Nature communications, 11(1), p. 3406.

Cristofanilli, M. et al. (2004) ‘Circulating tumor cells, disease progression, and survival in metastatic breast cancer’, The New England journal of medicine, 351(8), pp. 781–791.

Danila, D. C. et al. (2007) ‘Circulating tumor cell number and prognosis in progressive castration-resistant prostate cancer’, Clinical cancer research: an official journal of the American Association for Cancer Research, 13(23), pp. 7053–7058.

Dill, E. A. et al. (2018) ‘IDO expression in breast cancer: an assessment of 281 primary and metastatic cases with comparison to PD-L1’, *Modern pathology: an official journal of the United States and Canadian Academy of Pathology*, Inc, 31(10), pp. 1513–1522.

Ding, H. et al. (2019) ‘Biological process activity transformation of single cell gene expression for cross-species alignment’, Nature communications, 10(1), p. 4899.

Ding, J. et al. (2019) ‘Systematic comparative analysis of single cell RNA-sequencing methods’, BioRxiv. Available at: https://www.biorxiv.org/content/10.1101/632216v1.abstract.

Durante, M. A. et al. (2020) ‘Single-cell analysis reveals new evolutionary complexity in uveal melanoma’, Nature communications, 11(1), p. 496.

Ebright, R. Y. et al. (2020) ‘Deregulation of ribosomal protein expression and translation promotes breast cancer metastasis’, Science, 367(6485), pp. 1468–1473.

Eelen, G. et al. (2008) ‘Expression of the BRCA1-interacting protein Brip1/BACH1/FANCJ is driven by E2F and correlates with human breast cancer malignancy’, Oncogene, 27(30), pp. 4233–4241.

Farace, F. et al. (2011) ‘A direct comparison of CellSearch and ISET for circulating tumour-cell detection in patients with metastatic carcinomas’, British journal of cancer, 105(6), pp. 847–853.

Fard, M. M., Thonet, T. and Gaussier, E. (2020) ‘Deep k-Means: Jointly clustering with k-Means and learning representations’, Pattern Recognition Letters, pp. 185–192. doi: 10.1016/j.patrec.2020.07.028.

Ferreira, M. M., Ramani, V. C. and Jeffrey, S. S. (2016) ‘Circulating tumor cell technologies’, Molecular oncology, 10(3), pp. 374–394.

Follain, G. et al. (2018) ‘Hemodynamic Forces Tune the Arrest, Adhesion, and Extravasation of Circulating Tumor Cells’, Developmental cell, 45(1), pp. 33–52.e12.

Fu, T. et al. (2019) ‘DDL: Deep Dictionary Learning for Predictive Phenotyping’, in IJCAI, pp. 5857–5863.

Gabriel, M. T. et al. (2016) ‘Circulating Tumor Cells: A Review of Non–EpCAM-Based Approaches for Cell Enrichment and Isolation’, Clinical chemistry, 62(4), pp. 571–581.

Gajewski, T. F., Schreiber, H. and Fu, Y.-X. (2013) ‘Innate and adaptive immune cells in the tumor microenvironment’, Nature Immunology, pp. 1014–1022. doi: 10.1038/ni.2703.

Giuliano, M. et al. (2011) ‘Circulating tumor cells as prognostic and predictive markers in metastatic breast cancer patients receiving first-line systemic treatment’, Breast cancer research: BCR, 13(3), p. R67.

Guo, M. et al. (2015) ‘SINCERA: A Pipeline for Single-Cell RNA-Seq Profiling Analysis’, PLoS computational biology, 11(11), p. e1004575.

Guo, X. et al. (2017) ‘Deep Clustering with Convolutional Autoencoders’, in Neural Information Processing. Springer International Publishing, pp. 373–382.

Gupta, K. et al. (2020) ‘The Cellular basis of the loss of smell in 2019-nCoV-infected individuals’, Briefings in bioinformatics. doi: 10.1093/bib/bbaa168.

Habli, Z. et al. (2020) ‘Circulating Tumor Cell Detection Technologies and Clinical Utility: Challenges and Opportunities’, Cancers, 12(7). doi: 10.3390/cancers12071930.

Hänzelmann, S., Castelo, R. and Guinney, J. (2013) ‘GSVA: gene set variation analysis for microarray and RNA-seq data’, BMC bioinformatics, 14, p. 7.

Hao, Y. et al. (2021) ‘Integrated analysis of multimodal single-cell data’, Cell, 184(13), pp. 3573–3587.e29.

Hong, Y., Fang, F. and Zhang, Q. (2016) ‘Circulating tumor cell clusters: What we know and what we expect (Review)’, International Journal of Oncology, pp. 2206–2216. doi: 10.3892/ijo.2016.3747.

Howe, K. L. et al. (2021) ‘Ensembl 2021’, Nucleic acids research, 49(D1), pp. D884–D891.

Ignatiadis, M., Sotiriou, C. and Pantel, K. (2012) Minimal Residual Disease and Circulating Tumor Cells in Breast Cancer. Springer Science & Business Media.

Ingvarsson, S. (2001) ‘FHIT alterations in breast cancer’, Seminars in cancer biology, 11(5), pp. 361–366.

Iyer, A. et al. (2020) ‘Integrative Analysis and Machine Learning based Characterization of Single Circulating Tumor Cells’, Journal of clinical medicine research, 9(4). doi: 10.3390/jcm9041206.

Jiang, Y. et al. (2016) ‘Assessing intratumor heterogeneity and tracking longitudinal and spatial clonal evolutionary history by next-generation sequencing’, Proceedings of the National Academy of Sciences of the United States of America, 113(37), pp. E5528–37.

Jin, L. et al. (2014) ‘Pathway-based analysis tools for complex diseases: a review’, Genomics, proteomics & bioinformatics, 12(5), pp. 210–220.

Jin, X. et al. (2019) ‘ERα is required for suppressing OCT4-induced proliferation of breast cancer cells via DNMT1/ISL1/ERK axis’, Cell proliferation, 52(4), p. e12612.

Jordan, N. V. et al. (2016) ‘HER2 expression identifies dynamic functional states within circulating breast cancer cells’, Nature, 537(7618), pp. 102–106.

Kamal, M. et al. (2017) ‘Circulating Tumor Cells in Breast Cancer: A Potential Liquid Biopsy’, in Van Pham, P. (ed.) Breast Cancer. Rijeka: IntechOpen.

Kang, J. B. et al. (2020) ‘Efficient and precise single-cell reference atlas mapping with Symphony’, bioRxiv. Available at: https://www.biorxiv.org/content/10.1101/2020.11.18.389189v1.abstract.

Kanwar, N. et al. (2015) ‘Identification of genomic signatures in circulating tumor cells from breast cancer’, International journal of cancer. Journal international du cancer, 137(2), pp. 332–344.

Kim, C. et al. (2018) ‘Chemoresistance Evolution in Triple-Negative Breast Cancer Delineated by Single-Cell Sequencing’, Cell, 173(4), pp. 879–893.e13.

Kiselev, V. Y. et al. (2017) ‘SC3: consensus clustering of single-cell RNA-seq data’, Nature methods, 14(5), pp. 483–486.

Kiselev, V. Y., Andrews, T. S. and Hemberg, M. (2019a) ‘Challenges in unsupervised clustering of single-cell RNA-seq data’, Nature reviews. Genetics, 20(5), pp. 273–282.

Kiselev, V. Y., Andrews, T. S. and Hemberg, M. (2019b) ‘Publisher Correction: Challenges in unsupervised clustering of single-cell RNA-seq data’, Nature reviews. Genetics, 20(5), p. 310.

Kitamura, T. (2018) ‘A negative regulator of metastasis promoting macrophages’, J Emerg Crit Care Med. Available at: https://www.research.ed.ac.uk/portal/files/64401795/Editorial_Kitamura_v2.docx.

Koch, C. et al. (2020) ‘Characterization of circulating breast cancer cells with tumorigenic and metastatic capacity’, EMBO molecular medicine, 12(9), p. e11908.

Krämer, A. et al. (2014) ‘Causal analysis approaches in Ingenuity Pathway Analysis’, Bioinformatics , 30(·), pp. 523–530.

Krebs, M. G. et al. (2014) ‘Molecular analysis of circulating tumour cells—biology and biomarkers’, Nature reviews. Clinical oncology, 11(3), pp. 129–144.

Kwa, M. and Esteva, F. J. (2018) ‘Detection and Clinical Implications of Occult Systemic Micrometastatic Breast Cancer’, The Breast, pp. 858–866.e3. doi: 10.1016/b978-0-323-35955-9.00066-0.

Law, C. W. et al. (2014) ‘voom: Precision weights unlock linear model analysis tools for RNA-seq read counts’, Genome biology, 15(2), p. R29.

Lee, Y., Guan, G. and Bhagat, A. A. (2018) ‘ClearCell® FX, a label-free microfluidics technology for enrichment of viable circulating tumor cells’, Cytometry. Part A: the journal of the International Society for Analytical Cytology, 93(12), pp. 1251–1254.

Li, B. and Dewey, C. N. (2011) ‘RSEM: accurate transcript quantification from RNA-Seq data with or without a reference genome’, BMC bioinformatics, 12, p. 323.

Li, H. et al. (2017) ‘Reference component analysis of single-cell transcriptomes elucidates cellular heterogeneity in human colorectal tumors’, Nature genetics, 49(5), pp. 708–718.

Ling, Y. et al. (2015) ‘Loss of heterozygosity in thyroid hormone receptor beta in invasive breast cancer’, Tumori, 101(5), pp. 572–577.

Liu, S. et al. (2020) ‘SOD1 Promotes Cell Proliferation and Metastasis in Non-small Cell Lung Cancer via an miR-409-3p/SOD1/SETDB1 Epigenetic Regulatory Feedforward Loop’, Frontiers in cell and developmental biology, 8, p. 213.

Liu, Y. and Cao, X. (2016) ‘Immunosuppressive cells in tumor immune escape and metastasis’, Journal of molecular medicine , 94(5), pp. 509–522.

Lobo, I. (2008) ‘Chromosome Abnormalities and Cancer Genetics’.

Macosko, E. Z. et al. (2015) ‘Highly Parallel Genome-wide Expression Profiling of Individual Cells Using Nanoliter Droplets’, Cell, 161(5), pp. 1202–1214.

Mafficini, A. and Scarpa, A. (2018) ‘Genomic landscape of pancreatic neuroendocrine tumours: the International Cancer Genome Consortium’, The Journal of endocrinology, 236(3), pp. R161–R167.

Mahdizadehaghdam, S. et al. (2019) ‘Deep Dictionary Learning: A PARametric NETwork Approach’, IEEE transactions on image processing: a publication of the IEEE Signal Processing Society. doi: 10.1109/TIP.2019.2914376.

Mathers, C. D. (2020) ‘History of global burden of disease assessment at the World Health Organization’, Archives of public health = Archives belges de sante publique, 78, p. 77.

McAllister, S. S. and Weinberg, R. A. (2014) ‘The tumour-induced systemic environment as a critical regulator of cancer progression and metastasis’, Nature cell biology, 16(8), pp. 717–727.

Mikolajczyk, S. D. et al. (2011) ‘Detection of EpCAM-Negative and Cytokeratin-Negative Circulating Tumor Cells in Peripheral Blood’, Journal of oncology, 2011, p. 252361.

Miller, M. C., Doyle, G. V. and Terstappen, L. W. M. M. (2010) ‘Significance of Circulating Tumor Cells Detected by the CellSearch System in Patients with Metastatic Breast Colorectal and Prostate Cancer’, Journal of oncology, 2010, p. 617421.

Nagrath, S. et al. (2007) ‘Isolation of rare circulating tumour cells in cancer patients by microchip technology’, Nature, 450(7173), pp. 1235–1239.

Natrajan, R. et al. (2012) ‘Functional characterization of the 19q12 amplicon in grade III breast cancers’, Breast cancer research: BCR, 14(2), p. R53.

Orsetti, B. et al. (2006) ‘Genetic profiling of chromosome 1 in breast cancer: mapping of regions of gains and losses and identification of candidate genes on 1q’, British journal of cancer, 95(10), pp. 1439–1447.

Ozkumur, E. et al. (2013) ‘Inertial focusing for tumor antigen-dependent and -independent sorting of rare circulating tumor cells’, Science translational medicine, 5(179), p. 179ra47.

Peng, X. et al. (2016) ‘Deep Subspace Clustering with Sparsity Prior’, in IJCAI, pp. 1925–1931.

Privitera, A. P., Barresi, V. and Condorelli, D. F. (2021) ‘Aberrations of Chromosomes 1 and 16 in Breast Cancer: A Framework for Cooperation of Transcriptionally Dysregulated Genes’, Cancers, 13(7). doi: 10.3390/cancers13071585.

Pucci, F. et al. (2016) ‘PF4 Promotes Platelet Production and Lung Cancer Growth’, Cell reports, 17(7), pp. 1764–1772.

Rack, B. et al. (2014) ‘Circulating tumor cells predict survival in early average-to-high risk breast cancer patients’, Journal of the National Cancer Institute, 106(5). doi: 10.1093/jnci/dju066.

Ramalingam, N. et al. (2017) ‘Corrigendum: Fluidic Logic Used in a Systems Approach to Enable Integrated Single-Cell Functional Analysis’, Frontiers in Bioengineering and Biotechnology. doi: 10.3389/fbioe.2017.00011.

Ramirez, A. K. et al. (2020) ‘Single-cell transcriptional networks in differentiating preadipocytes suggest drivers associated with tissue heterogeneity’, Nature communications, 11(1), p. 2117.

Ranjan, B. et al. (2021) ‘scConsensus: combining supervised and unsupervised clustering for cell type identification in single-cell RNA sequencing data’, BMC bioinformatics, 22(1), p. 186.

Riethdorf, S. et al. (2007) ‘Detection of circulating tumor cells in peripheral blood of patients with metastatic breast cancer: a validation study of the CellSearch system’, Clinical cancer research: an official journal of the American Association for Cancer Research, 13(3), pp. 920–928.

Ritchie, M. E. et al. (2015) ‘limma powers differential expression analyses for RNA-sequencing and microarray studies’, Nucleic acids research, 43(7), p. e47.

Robinson, M. D., McCarthy, D. J. and Smyth, G. K. (2010) ‘edgeR: a Bioconductor package for differential expression analysis of digital gene expression data’, Bioinformatics , 26(1), pp. 139–140.

Sarioglu, A. F. et al. (2015) ‘A microfluidic device for label-free, physical capture of circulating tumor cell clusters’, Nature methods, 12(7), pp. 685–691.

Senchenko, V. N. et al. (2004) ‘Discovery of frequent homozygous deletions in chromosome 3p21.3 LUCA and AP20 regions in renal, lung and breast carcinomas’, Oncogene, 23(34), pp. 5719–5728.

Sheikhpour, E. et al. (2018) ‘A Survey on the Role of Interleukin-10 in Breast Cancer: A Narrative’, Reports of biochemistry & molecular biology, 7(1), pp. 30–37.

Shenoy, A. K. and Lu, J. (2016) ‘Cancer cells remodel themselves and vasculature to overcome the endothelial barrier’, Cancer letters, 380(2), pp. 534–544.

Siegel, R. L., Miller, K. D. and Jemal, A. (2015) ‘Cancer statistics, 2015’, CA: a cancer journal for clinicians, 65(1), pp. 5–29.

Singhal, V. and Majumdar, A. (2018) ‘Majorization Minimization Technique for Optimally Solving Deep Dictionary Learning’, Neural Processing Letters, pp. 799–814. doi: 10.1007/s11063-017-9603-9.

Sinha, D. et al. (2018) ‘dropClust: efficient clustering of ultra-large scRNA-seq data’, Nucleic acids research, 46(6), p. e36.

Sinha, D. et al. (2019) ‘Improved dropClust R package with integrative analysis support for scRNA-seq data’, Bioinformatics . doi: 10.1093/bioinformatics/btz823.

Stott, S. L. et al. (2010) ‘Isolation of circulating tumor cells using a microvortex-generating herringbone-chip’, Proceedings of the National Academy of Sciences of the United States of America, 107(43), pp. 18392–18397.

Stouffer, S. A. et al. (1949) ‘The American soldier: Adjustment during army life. (Studies in social psychology in World War II), Vol. 1’, 1, p. 599.

Subramanian, A. et al. (2005) ‘Gene set enrichment analysis: a knowledge-based approach for interpreting genome-wide expression profiles’, Proceedings of the National Academy of Sciences of the United States of America, 102(43), pp. 15545–15550.

Sudmant, P. H. et al. (2015) ‘An integrated map of structural variation in 2,504 human genomes’, Nature, 526(7571), pp. 75–81.

Sung, H. et al. (2021) ‘Global cancer statistics 2020: GLOBOCAN estimates of incidence and mortality worldwide for 36 cancers in 185 countries’, CA: a cancer journal for clinicians. Available at: https://acsjournals.onlinelibrary.wiley.com/doi/abs/10.3322/caac.21660.

Szczerba, B. M. et al. (2019) ‘Neutrophils escort circulating tumour cells to enable cell cycle progression’, Nature, 566(7745), pp. 553–557.

Tang, H. et al. (2020) ‘When Dictionary Learning Meets Deep Learning: Deep Dictionary Learning and Coding Network for Image Recognition With Limited Data’, *IEEE transactions on neural networks and learning systems*, PP. doi: 10.1109/TNNLS.2020.2997289.

Tariyal, S. et al. (2016) ‘Deep Dictionary Learning’, IEEE Access, 4, pp. 10096–10109.

Thery, L. et al. (2019) ‘Circulating Tumor Cells in Early Breast Cancer’, JNCI cancer spectrum, 3(2), p. kz026.

Tian, L. et al. (2019) ‘Benchmarking single cell RNA-sequencing analysis pipelines using mixture control experiments’, Nature methods, 16(6), pp. 479–487.

Tickle, T. et al. (2019) ‘inferCNV of the Trinity CTAT Project’, Klarman Cell Observatory, Broad Institute of MIT and Harvard.

Ting, D. T. et al. (2014) ‘Single-cell RNA sequencing identifies extracellular matrix gene expression by pancreatic circulating tumor cells’, Cell reports, 8(6), pp. 1905–1918.

Tirosh, I. et al. (2016) ‘Dissecting the multicellular ecosystem of metastatic melanoma by single-cell RNA-seq’, Science, 352(6282), pp. 189–196.

Tošić, I. and Frossard, P. (2011) ‘Dictionary Learning’, IEEE Signal Processing Magazine, 28(2), pp. 27–38.

Tsai, W.-S. et al. (2016) ‘Circulating Tumor Cell Count Correlates with Colorectal Neoplasm Progression and Is a Prognostic Marker for Distant Metastasis in Non-Metastatic Patients’, Scientific reports, 6, p. 24517.

Turner, N. et al. (2010) ‘Integrative molecular profiling of triple negative breast cancers identifies amplicon drivers and potential therapeutic targets’, Oncogene, 29(14), pp. 2013–2023.

Urrutia, E. et al. (2018) ‘Integrative pipeline for profiling DNA copy number and inferring tumor phylogeny’, Bioinformatics , 34(12), pp. 2126–2128.

Velten, L. et al. (2017) ‘Human haematopoietic stem cell lineage commitment is a continuous process’, Nature cell biology, 19(4), pp. 271–281.

Wang, L. et al. (2016) ‘Promise and limits of the CellSearch platform for evaluating pharmacodynamics in circulating tumor cells’, Seminars in oncology, 43(4), pp. 464–475.

Wang, S. et al. (2020) ‘Single-Cell Transcriptomic Atlas of Primate Ovarian Aging’, Cell, 180(3), pp. 585–600.e19.

Ward, M. P. et al. (2021) ‘Platelets, immune cells and the coagulation cascade; friend or foe of the circulating tumour cell?’, Molecular cancer, 20(1), p. 59.

Warkiani, M. E. et al. (2014) ‘Slanted spiral microfluidics for the ultra-fast, label-free isolation of circulating tumor cells’, Lab on a chip, 14(1), pp. 128–137.

Weinstein, J. N. et al. (2013) ‘The Cancer Genome Atlas Pan-Cancer analysis project’, Nature Genetics, pp. 1113–1120. doi: 10.1038/ng.2764.

Wolf, F. A., Angerer, P. and Theis, F. J. (2018) ‘SCANPY: large-scale single-cell gene expression data analysis’, Genome biology, 19(1), p. 15.

Xie, J., Girshick, R. and Farhadi, A. (2016) ‘Unsupervised Deep Embedding for Clustering Analysis’, in Balcan, M. F. and Weinberger, K. Q. (eds) Proceedings of The 33rd International Conference on Machine Learning. New York, New York, USA: PMLR (Proceedings of Machine Learning Research), pp. 478–487.

Xu, L. et al. (2015) ‘Optimization and Evaluation of a Novel Size Based Circulating Tumor Cell Isolation System’, PloS one, 10(9), p. e0138032.

Yang, B. et al. (2017) ‘Towards K-means-friendly Spaces: Simultaneous Deep Learning and Clustering’, in Precup, D. and Teh, Y. W. (eds) Proceedings of the 34th International Conference on Machine Learning. International Convention Centre, Sydney, Australia: PMLR (Proceedings of Machine Learning Research), pp. 3861–3870.

Yang, X. et al. (2019) ‘Deep spectral clustering using dual autoencoder network’, in *Proceedings of the IEEE/CVF Conference on Computer Vision and Pattern Recognition*, pp. 4066–4075.

Yip, S. H. et al. (2017) ‘Linnorm: improved statistical analysis for single cell RNA-seq expression data’, Nucleic acids research, 45(22), p. e179.

Yu, M. et al. (2014) ‘Cancer therapy. Ex vivo culture of circulating breast tumor cells for individualized testing of drug susceptibility’, Science, 345(6193), pp. 216–220.

Zhang, H. et al. (2021) ‘Detection Methods and Clinical Applications of Circulating Tumor Cells in Breast Cancer’, Frontiers in oncology, 11, p. 652253.

Zhao, W. et al. (2017) ‘Decreasing Eukaryotic Initiation Factor 3C (EIF3C) Suppresses Proliferation and Stimulates Apoptosis in Breast Cancer Cell Lines Through Mammalian Target of Rapamycin (mTOR) Pathway’, Medical science monitor: international medical journal of experimental and clinical research, 23, pp. 4182–4191.

Zheng, Y. et al. (2017) ‘Expression of β-globin by cancer cells promotes cell survival during blood-borne dissemination’, Nature communications, 8, p. 14344.

Zhou, Y. et al. (2020) ‘Single-cell RNA landscape of intratumoral heterogeneity and immunosuppressive microenvironment in advanced osteosarcoma’, Nature communications, 11(1), p. 6322.

