## Supplemental Data 1 for "Marker-free characterization of single live circulating tumor cell full-length transcriptomes"

**Supplementary Table S1.** Description of CTCs enriched by ClearCell^®^ FX and Polaris^TM^ workflow

| **Patient** | **#CTCs** | **Hormone Receptor Status** |
| --- | --- | --- |
| **P1** | 11 | ER-/PR-/HER2- |
| **P3** | 13 | ER+/PR+/HER2- |
| **P4** | 12 | ER+/PR+/HER2- |
| **P5** | 32 | ER+/PR+/HER2- |
| **P7** | 5 | ER-/PR-/HER2+ |
| **P9** | 8 | ER-/PR-/HER2+ |
| **Total** | **81** |  |
