## Supplemental Data 2 for "Marker-free characterization of single live circulating tumor cell full-length transcriptomes"

**Supplementary Table S2.** Summary of CTCs enriched through ClearCell^®^ FX-Polaris^TM^ workflow.

| **Patient** | **#CTC** | **Hormone Receptor Status** |
| --- | --- | --- |
| **P1** | 11 | ER-/PR-/HER2- |
| **P3** | 10 | ER+/PR+/HER2- |
| **P4** | 12 | ER+/PR+/HER2- |
| **P5** | 32 | ER+/PR+/HER2- |
| **P7** | 4 | ER-/PR-/HER2+ |
| **P9** | 3 | ER-/PR-/HER2+ |
| **Total** | **72** |  |
