## Supplementary Table S3: Description of datasets. for "Marker-free characterization of single live circulating tumor cell full-length transcriptomes"

| **Study** | **Dataset** | **#WBCs** | **#CTCs** |
| --- | --- | --- | --- |
| **Study #1** | **Velten et al. ( GSE75478 )** (Velten *et al.*, 2017) | 1035 HSCs and progenitors | 0 |
|  | **Ting et al. (GSE60407)** (Ting *et al.*, 2014) | 0 | 7 pancreatic CTCs |
|  | **Yu et al. (GSE55807)**  (Yu *et al.*, 2014) | 0 | 6 breast CTCs |
|  | **Sarioglu et al. (GSE67939)**  (Sarioglu *et al.*, 2015) | 2 WBCs | 15 breast CTCs |
|  | **Aceto et al. (GSE51827)**  (Aceto *et al.*, 2014) | 0 | 29 breast CTCs and CTC clusters |
|  | **Zheng et al. (GSE74639)**  (Zheng *et al.*, 2017) | 0 | 10 Lung CTCs  6 single cells from Primary Tumor |
|  | **Jordan et al. (GSE75367)**  (Jordan *et al.*, 2016) | 0 | 74 breast CTCs |
| **Study #2** | **Ebright et al.** (GSE144494)  (Ebright *et al.*, 2020) | 0 | 824 breast CTCs |
|  | Poonia et al. (ClearCell^®^ FX Polaris^TM^) | 0 | 81 CTCs breast cancer |
|  | **Ding et al. WBC1** (<https://portals.broadinstitute.org/single_cell>)  (Ding *et al.*, 2019) | 376 | 0 |
|  | **Ding et al. WBC1** (<https://portals.broadinstitute.org/single_cell>)  (Ding *et al.*, 2019) | 376 | 0 |
