## Supplementary Table S5: Cluster wise differentially upregulated genes. for "Marker-free characterization of single live circulating tumor cell full-length transcriptomes"

| **Custer_0** | **Custer_1** | **Custer_2** | **Custer_3** |
| --- | --- | --- | --- |
| MTRNR2L8 | ACRBP | ING5 | HBB |
| TMSB4X | TUBB1 | HBB | KRT18 |
| RPS28 | CLEC1B | MAP1LC3C | AGR2 |
| RPL13 | SELP | EXD1 | RPA3 |
| CD52 | PPBP | SLC36A2 | KRT19 |
| RPS9 | TSC22D1 | ITGA2B | TSR3 |
| MTRNR2L1 | SPARC | FBLIM1 | CLTB |
| CORO1A | RUFY1 | SP100 | ELF3 |
| RPS18 | ITGA2B | VSTM4 | MLPH |
| RPS15 | MPP1 | ZFP30 | TUBB6 |
| RPL28 | PF4 | ITIH5 | DHRS4 |
| RPL3 | ALOX12 | LMOD3 | GRHPR |
| RPL19 | TMEM40 | GATAD1 | DHRS4L2 |
| ITGB2 | HBB | AS3MT | FXYD3 |
| RPS17 | GSTO1 | COX6B2 | VPS51 |
| RPL9 | C2orf88 | C1orf229 | MGP |
| RPL10A | RGS10 | QPCTL | GPKOW |
| RPL23A | PDLIM1 | HSD17B13 | XBP1 |
| RPL17 | VCL | ZNF677 | MUC1 |
| RPS23 | TSPAN33 | GPR82 | HSPB1 |
| MALAT1 | NFE2 | MTFMT | GRINA |
| RPS29 | DAB2 | IDO1 | TACSTD2 |
| RPL26 | XPNPEP1 | METTL2B | BAIAP2 |
| RPLP2 | TMEM140 | LYRM4 | EIF3C |
| RAC2 | ODC1 | TMEM213 | MYOF |
| RPS26 | ITGB5 | MED18 | HSP90AB1 |
| RPS27 | GNG11 | FCAR | RPS2 |
| MTRNR2L6 | TUBA4A | TAF8 | EPCAM |
| EEF1A1 | TREML1 | C12orf65 | BANF1 |
| RPS3A | PTGS1 | NMNAT1 | PRDX2 |
| CD53 | ESAM | APOBEC3A | AZGP1 |
| RPS16 | ABCC3 | AP1S3 | KPNA2 |
| RPL18A | SMOX | ZNF250 | HSP90AA1 |
| LSP1 | NRGN | GDPD1 | BLVRB |
| RPS25 | CLU | AKIP1 | S100A14 |
| RPL18 | RGS18 | MOG | CCT3 |
| RPL6 | GFI1B | CXorf56 | DYNLL1 |
| RPL29 | TAL1 | DAND5 | DDR1 |
| RPL5 | TLN1 | NEK2 | TRIM28 |
| RPS3 | MMD | BPNT1 | CD81 |
| HNRNPA1 | ICAM2 | ENTPD4 | S100P |
| RPS14 | GRAP2 | APOL1 | PHB |
| RPS5 | CDKN2D | PSTPIP2 | KRT8 |
| CXCR4 | RDH11 | AIPL1 | PEBP1 |
| RPL32 | PLEKHO1 | PECR | RAB24 |
| RPL34 | CDKN1A | ENTPD1 | NME1 |
| RPS4X | RAB11A | TRMT10B | JTB |
| RPS13 | CTSA | ZBTB3 | CAPNS1 |
| RPL13A | RAB32 | HCAR1 | NDUFB9 |
| RPS6 | CTDSPL | VSIG1 | HDGF |
| RPL7 | ETFA | ZNF483 | IFI27 |
| RPS15A | PRKAR2B | ZNF554 | S100A11 |
| RPL10 | GPX1 | GINS4 | CYC1 |
| RPS19 | ENDOD1 | TRAF3IP2 | SOD1 |
| RPL7A | VSIG2 | SHROOM4 | PPA1 |
| RPL11 | MFSD1 | CC2D2A | FDPS |
| RPL39 | PTCRA | MANEAL | PSMB7 |
| IL2RG | LIMS1 | C15orf40 | SPINT2 |
| RPL31 | ZNF185 | ZFP42 | RPF1 |
| RPSA | TPM4 | GREB1 | DSTN |
| RPL8 | MEPCE | SKA1 | RPLP0 |
| RPL14 | MAX | LNX2 | ANXA2 |
| RPL37A | LGALSL | CHRNB1 | BAG6 |
| CD48 | GP6 | NFKBIB | PSMA4 |
| RPL21 | F13A1 | INIP | TXN |
| RPL30 | TPST2 | SEC14L4 | HDLBP |
| LCP1 | MYLK | TRPV1 | ATP6V0C |
| RPS10 | THBS1 | VPS33A | HSPD1 |
| EEF1G | TSPAN18 | SCD5 | RNF187 |
| RPS7 | CTTN | LRRN4CL | TPI1 |
| PTPRC | NEXN | THAP6 | PSMB4 |
| LAPTM5 | GAS2L1 | SLC12A8 | SNRPE |
| HNRNPH1 | TIMP1 | CYP1A2 | RAB25 |
| EEF1D | BIN2 | PDLIM5 | ALAS2 |
| VIM | BCL2L1 | ASTN2 | PSMD4 |
| FAU | PGRMC1 | ZNF527 | DCTPP1 |
| S100A4 | P2RX1 | CCL22 | UQCRQ |
| RPLP1 | NCOA4 | SPTLC1 | HSPA8 |
| PTPRCAP | EHD3 | ZBTB8A | TSPAN13 |
| TXNIP | LAT | DYDC1 | STMN1 |
| IFITM1 | RAP1B | ABCB5 | CDK4 |
| SLC25A6 | LTBP1 | ZNF620 | HBD |
| CALM1 | PTPN18 | ADRA1A | MDK |
| RPL27 | ILK | FAM74A1 | EIF6 |
| C6orf62 | PEAR1 | SHOX | SPINT1 |
| RPL36A | TUBA8 | PLCXD1 | KRT7 |
| RPL36 | FAM110A | SLC28A2 | LDHA |
| CD37 | TALDO1 | BRIP1 | CD63 |
| IL32 | UBL4A | MAB21L3 | TUBB4B |
| RPS12 | SLC2A3 | ZNF850 | PLIN3 |
| CD74 | RAB31 | SEMA3E | RAN |
| RPL12 | LAMTOR1 | ARGFX | PSMB2 |
| RPS21 | VDAC3 | CRX | CHCHD2 |
| NPM1 | DIAPH1 | CYP4V2 | PSMB5 |
| EEF1B2 | EIF2AK1 | WDR92 | SELENBP1 |
| IFITM2 | SNCA | PCBD2 | NQO1 |
| RPS11 | TUBA1C | TNFAIP8L1 | LGALS3BP |
| RPL35 | GMPR | RHD | TYMS |
| RPS24 | DNM3 | CABP4 | BSCL2 |
| RPS8 | MFAP3L | TBXA2R | AP2M1 |
| SRSF5 | PLA2G12A | DGKB | SLC1A5 |
| RPL41 | MEIS1 | NRIP3 | PRDX1 |
| NKG7 | GNAQ | CYP51A1 | CCT2 |
| NACA | TTC7B | MOCS3 | NDUFS2 |
| HCLS1 | LGALS12 | PPARD | COX6C |
| RPL27A | PDLIM7 | ZNF669 | HBA2 |
| DAZAP2 | GLA | SLC14A2 | BLVRA |
| RPLP0 | ZYX | MEFV | PSMC5 |
| PABPC1 | PDZK1IP1 | RAB42 | TFF1 |
| MTRNR2L10 | PCSK6 | CCBE1 | AMZ2 |
| ANXA1 | CNST | QPRT | TMEM205 |
| PCSK5 | MKRN1 | SLC31A1 | HSPA1A |
| RPL38 | STOM | TOR1AIP2 | CCT5 |
| RPL35A | THOC2 | PRR11 | REPIN1 |
| RPS27A | PTTG1IP | AP4S1 | PDCD6 |
| RPL37 | PTP4A2 | LRRC2 | DUS1L |
| RPL24 | WRNIP1 | LRRC27 | ERGIC3 |
| TMSB10 | RAB27B | MILR1 | SLC25A3 |
| EVI2B | SAT1 | C21orf62 | C19orf53 |
| RPL23 | ITGB3 | ATCAY | CDK5RAP3 |
| RBM3 | GNAZ | ZNF490 | PTMS |
| PFDN5 | GATA1 | PGAM5 | DYNLRB1 |
| MORC4 | CPNE5 | TSTD3 | PSME2 |
| SELPLG | CCND3 | ASB11 | NADK |
| ZFP36L1 | PNP | HTRA4 | RPL7 |
| NCL | GP1BA | CCDC142 | TCEAL4 |
| RPL4 | PTPN12 | SULT2A1 | XRCC5 |
| TRAF3IP3 | TACC3 | CPM | CRABP2 |
| ASB3 | NAP1L1 | SLC50A1 | BTF3 |
| DDX5 | C1orf198 | CDH23 | UTP14A |
| LIMD2 | MGLL | C1orf210 | MDH2 |
| PPIA | TJP2 | ZNF835 | LRRC59 |
| ZAP70 | GRK5 | EXPH5 | ECHS1 |
| CDC42 | IFRD1 | CCDC170 | MRPL51 |
| ARHGAP15 | GPX4 | TBCCD1 | PSMD1 |
| RPS20 | INSIG1 | PIWIL2 | NOP56 |
| CST7 | SAV1 | EPHA10 | PRDX4 |
| SELL | R3HDM4 | TMEM19 | SEPHS2 |
| RPL15 | GNAS | FADS6 | EFNA1 |
| LITAF | PLEK | SBSPON | MRPS26 |
| TOMM7 | CALM3 | ZNF793 | PSMA7 |
| HNRNPC | PDE5A | OPA3 | SNRPC |
| PPP2R5C | TMEM164 | KCNA7 | TSPAN1 |
| PFN1 | CD9 | RNF170 | ILF2 |
| TPT1 | FERMT3 | FBXO27 | SNF8 |
| PLAC8 | SIAH2 | GGT6 | SLC25A5 |
| CNBP | PNKD | ZNF526 | PRSS8 |
| HNRNPK | DIMT1 | LRTOMT | MRFAP1 |
| AHNAK | STK24 | ZNF430 | AGR3 |
| GNG2 | CLCN3 | HSCB | SPDEF |
| RPL22 | AMD1 | L2HGDH | TOP2A |
| EML4 | SUSD1 | SUV39H2 | SNRNP25 |
| TRIM22 | UBE2E3 | SLC9A7 | NHP2 |
| S1PR1 | PDK1 | CD82 | PPP1R14B |
| PRF1 | ATF4 | METTL21A | PCBP1 |
| CD3D | UBE2F | PDE6A | PSMD8 |
| ZFP36 | WDR1 | NLRP12 | TUFM |
| COX4I1 | SYTL4 | SAA2 | TCP1 |
| TNFAIP3 | KCTD10 | HIF3A | SLC9A3R1 |
| ARHGDIB | CMIP | TMEM130 | RAC1 |
| BTF3 | TNFSF4 | MBOAT1 | GLO1 |
| SUMO2 | ABHD4 | FBXO6 | S100A16 |
| PTMA | ITM2B | TATDN3 | AIDA |
| STAT4 | SLA2 | ADIPOQ | FBL |
| TRA2A | SERPINB1 | SLC35E3 | PSMC3 |
| CD44 | KIF2A | AGMAT | SMAP1 |
| CNN2 | EMC3 | POU5F1 | RAB13 |
| DUSP1 | MICU1 | AFMID | RPSA |
| CFL1 | PPM1A | KCNJ3 | COX6A1 |
| IL10RA | PKM | BHMT2 | CLTC |
| TMEM45A | TRIM58 | RBPMS2 | CHMP2A |
| INPP5D | SNN | SLC52A1 | HSD17B10 |
| HSPA8 | CD63 | DMC1 | ZG16B |
| LCK | ANTXR2 | PBOV1 | SLC25A39 |
| CYTIP | PARD3 | ZNF506 | BSG |
| ANKRD44 | FYN | FBXL13 | AHCY |
| HCST | PRKCB | ARSK | SEC16A |
| SMG1 | BNIP3L | FUT1 | TUBB |
| CAPZA1 | PDCD10 | STAC2 | PPDPF |
| STK17B | MMRN1 | NUP43 | DBI |
| GIMAP4 | BMP6 | OLAH | RPL8 |
| EIF3F | PIP4K2A | IL10 | EPHX1 |
| S100A10 | CCNG1 | ACBD7 | TRIB1 |
| GIMAP7 | CHD1L | WNT7B | CFB |
| ARPC3 | CMTM5 | PGM5P2 | TAF9 |
| TBC1D10C | CD99 | SIX4 | SERPINA3 |
| SASH3 | RNF11 | SNHG7 | ESRP1 |
| CDC42SE1 | PRDX6 | TMEM105 | CTNNA1 |
| DDX6 | RAB37 | SLC25A15 | SRM |
| UBA52 | RGS6 | FAM161A | BST2 |
| EIF2S3 | PITPNM2 | ZSCAN2 | WDR34 |
| FOS | TBPL1 | BCL2L15 | FKBP4 |
| GZMA | CENPT | METTL6 | SMARCA4 |
| PSAP | CCDC92 | TNFAIP8L3 | CA12 |
| UCP2 | TAX1BP3 | KREMEN1 | VDAC2 |
| FTL | RSU1 | B3GALNT2 | NDUFA4 |
| EZR | SEC14L1 | C8orf86 | GATA3 |
| TAPBP | DAPP1 | ZNF865 | ANXA11 |
| HMGN2 | SQSTM1 | SLC7A14 | ERBB3 |
| FLNA | SPOCD1 | PTCHD4 | RPL12 |
