## Supplementary Table S7: Differential genes across CTCs of three different subtypes. for "Marker-free characterization of single live circulating tumor cell full-length transcriptomes"

| **Gene name** | **Subtype** |
| --- | --- |
| JUND | ER+/PR+ |
| CLEC4GP1 | ER+/PR+ |
| ATF3 | ER+/PR+ |
| DNAJC21 | ER+/PR+ |
| DLX6-AS1 | ER+/PR+ |
| SLC25A37 | ER+/PR+ |
| SBSPON | ER+/PR+ |
| ZEB2 | ER+/PR+ |
| PDE11A | ER+/PR+ |
| MYADML | ER+/PR+ |
| C12orf29 | TNBC |
| PPEF1 | TNBC |
| CISD1 | TNBC |
| EIF3C | TNBC |
| CDC42 | TNBC |
| GNAQ | TNBC |
| FOXP1 | TNBC |
| ACTR3 | TNBC |
| DEFB107A | TNBC |
| DEFB107B | TNBC |
| RAD54L2 | HER2+ |
| DIS3L2 | HER2+ |
| FLT4 | HER2+ |
| HMGN4 | HER2+ |
| EIF4E | HER2+ |
| CMC2 | HER2+ |
| PTPRC | HER2+ |
| MAP3K15 | HER2+ |
| CACYBP | HER2+ |
| NPAS3 | HER2+ |
