## Supplementary Table S8: CNV events in CTCs. for "Marker-free characterization of single live circulating tumor cell full-length transcriptomes"

| **Class** | **State** | **Chromosome arm** |
| --- | --- | --- |
| Poonia et al. | 2 | chr1_p36.33 |
| Poonia et al. | 4 | chr1_p36.32 |
| Poonia et al. | 4 | chr1_p35.2 |
| Poonia et al. | 2 | chr1_q23.1 |
| Poonia et al. | 4 | chr1_q25.3 |
| Poonia et al. | 4 | chr1_q32.1 |
| Poonia et al. | 5 | chr1_q32.3 |
| Poonia et al. | 5 | chr1_q41 |
| Poonia et al. | 2 | chr1_q42.13 |
| Poonia et al. | 4 | chr1_q42.3 |
| Poonia et al. | 5 | chr1_q44 |
| Poonia et al. | 2 | chr2_p23.3 |
| Poonia et al. | 2 | chr2_p13.3 |
| Poonia et al. | 2 | chr2_q24.1 |
| Poonia et al. | 2 | chr2_q32.3 |
| Poonia et al. | 4 | chr2_q33.1 |
| Poonia et al. | 5 | chr2_q34 |
| Poonia et al. | 4 | chr2_q35 |
| Poonia et al. | 2 | chr2_q37.3 |
| Poonia et al. | 2 | chr3_p24.3 |
| Poonia et al. | 4 | chr3_p22.1 |
| Poonia et al. | 2 | chr3_p21.31 |
| Poonia et al. | 4 | chr3_q21.2 |
| Poonia et al. | 4 | chr3_q25.32 |
| Poonia et al. | 5 | chr3_q26.1 |
| Poonia et al. | 6 | chr3_q27.1 |
| Poonia et al. | 5 | chr3_q29 |
| Poonia et al. | 4 | chr4_p15.32 |
| Poonia et al. | 5 | chr4_p15.1 |
| Poonia et al. | 4 | chr4_q22.1 |
| Poonia et al. | 4 | chr4_q34.2 |
| Poonia et al. | 4 | chr4_q35.2 |
| Poonia et al. | 4 | chr5_p15.33 |
| Poonia et al. | 4 | chr5_q12.3 |
| Poonia et al. | 5 | chr5_q14.1 |
| Poonia et al. | 4 | chr5_q14.2 |
| Poonia et al. | 4 | chr5_q31.1 |
| Poonia et al. | 2 | chr5_q35.3 |
| Poonia et al. | 4 | chr6_p25.3 |
| Poonia et al. | 2 | chr6_p21.33 |
| Poonia et al. | 4 | chr6_p21.31 |
| Poonia et al. | 4 | chr6_q12 |
| Poonia et al. | 4 | chr6_q21 |
| Poonia et al. | 4 | chr6_q22.31 |
| Poonia et al. | 4 | chr6_q25.3 |
| Poonia et al. | 2 | chr6_q27 |
| Poonia et al. | 4 | chr7_q22.1 |
| Poonia et al. | 4 | chr7_q31.1 |
| Poonia et al. | 5 | chr7_q32.2 |
| Poonia et al. | 4 | chr7_q34 |
| Poonia et al. | 5 | chr8_p23.3 |
| Poonia et al. | 4 | chr8_p23.1 |
| Poonia et al. | 4 | chr8_q21.3 |
| Poonia et al. | 2 | chr8_q24.13 |
| Poonia et al. | 4 | chr9_q31.2 |
| Poonia et al. | 5 | chr9_q33.2 |
| Poonia et al. | 4 | chr9_q33.3 |
| Poonia et al. | 2 | chr9_q34.3 |
| Poonia et al. | 4 | chr10_p15.1 |
| Poonia et al. | 5 | chr10_p13 |
| Poonia et al. | 4 | chr10_p12.1 |
| Poonia et al. | 4 | chr10_q24.1 |
| Poonia et al. | 2 | chr11_p15.5 |
| Poonia et al. | 3 | chr11_p15.1 |
| Poonia et al. | 2 | chr11_q14.1 |
| Poonia et al. | 4 | chr11_q23.1 |
| Poonia et al. | 4 | chr11_q24.2 |
| Poonia et al. | 5 | chr11_q25 |
| Poonia et al. | 2 | chr12_p13.33 |
| Poonia et al. | 4 | chr12_p12.3 |
| Poonia et al. | 5 | chr12_q12 |
| Poonia et al. | 4 | chr12_q13.11 |
| Poonia et al. | 2 | chr12_q13.3 |
| Poonia et al. | 4 | chr12_q14.2 |
| Poonia et al. | 5 | chr12_q15 |
| Poonia et al. | 4 | chr12_q21.1 |
| Poonia et al. | 4 | chr12_q23.3 |
| Poonia et al. | 2 | chr12_q24.31 |
| Poonia et al. | 2 | chr13_q34 |
| Poonia et al. | 2 | chr14_q22.1 |
| Poonia et al. | 2 | chr14_q24.1 |
| Poonia et al. | 2 | chr14_q32.33 |
| Poonia et al. | 4 | chr15_q14 |
| Poonia et al. | 5 | chr15_q15.3 |
| Poonia et al. | 4 | chr15_q21.3 |
| Poonia et al. | 2 | chr15_q26.1 |
| Poonia et al. | 2 | chr16_p13.3 |
| Poonia et al. | 2 | chr16_p12.1 |
| Poonia et al. | 4 | chr16_q12.1 |
| Poonia et al. | 2 | chr16_q22.1 |
| Poonia et al. | 5 | chr16_q23.1 |
| Poonia et al. | 2 | chr16_q24.3 |
| Poonia et al. | 5 | chr17_p13.3 |
| Poonia et al. | 2 | chr17_p13.2 |
| Poonia et al. | 4 | chr17_q11.2 |
| Poonia et al. | 2 | chr17_q22 |
| Poonia et al. | 2 | chr17_q25.3 |
| Poonia et al. | 2 | chr19_p13.3 |
| Poonia et al. | 4 | chr19_p13.11 |
| Poonia et al. | 6 | chr19_q12 |
| Poonia et al. | 5 | chr19_q13.11 |
| Poonia et al. | 2 | chr19_q13.33 |
| Poonia et al. | 5 | chr19_q13.41 |
| Poonia et al. | 6 | chr19_q13.42 |
| Poonia et al. | 6 | chr19_q13.43 |
| Poonia et al. | 2 | chr20_q11.23 |
| Poonia et al. | 4 | chr20_q13.33 |
| Ebright et al. | 2 | chr1_p22.2 |
| Ebright et al. | 2 | chr2_q12.1 |
| Ebright et al. | 3 | chr2_q36.1 |
| Ebright et al. | 2 | chr11_q23.3 |
| Ebright et al. | 2 | chr19_p13.2 |
| Ebright et al. | 3 | chr20_p13 |
